# Untangling the Indonesian tangle net fishery: describing a data-poor fishery targeting large threatened rays (Order Batoidea)

**DOI:** 10.1101/608935

**Authors:** Brooke M. D’Alberto, William T. White, Andrew Chin, Dharmadi, Colin A. Simpfendorfer

## Abstract

1. Shark-like rays (Order Rhinopristiformes) are among the most threatened families of marine fish. Yet little is known about their populations, as these rays are normally taken as opportunistic catch in fisheries targeting other species and are thus poorly reported. One exception is the Indonesian tangle net fishery, which targets shark-like rays.
2. Market surveys of Muara Angke landing site in Jakarta, north-western Java (including one frozen shipment from Benoa Harbour, Bali), were conducted between 2001 and 2005, and recorded landed catch for this fishery. Recent catch data from Indonesian Capture Fisheries (2017 – 2018) were also examined to provide contemporary information about landed catch.
3. 1,559 elasmobranchs (sharks and rays) were recorded, comprised of 24 species of rays and nine species of sharks. The most abundant species landed were the pink whipray *Pateobatis fai* and the bottlenose wedgefish *Rhynchobatus australiae*, the latter being the main target species.
4. Catch composition varied based on differences in species catchability and may also be indicative of localized declines. The fishery was highly selective for larger sized individuals, however smaller size classes of target species were also caught in other Indonesian fisheries resulting in fishing pressure across all age classes.
5. Evidence of substantial declines in global landings of wedgefish species, and the observed shift in catch composition in the Indonesian tangle net fishery, increases concerns about the status of shark-like rays and stingrays in Indonesia.

## 1. Introduction

Rays (Superorder Batoidae) are among the most threatened groups of chondrichthyans (sharks, rays, and chimaeras) (Dulvy et al., 2014a). Substantial declines in populations, catch rates and landings, as well as localised extinctions have been reported for several ray species (Moore, 2015). Declines in elasmobranchs populations are mainly the result of the rapid expansion of chondrichthyan catch in target and non-target fisheries (Dulvy et al., 2014a, Clarke et al., 2006), and the globalisation of trade (Lack and Sant, 2009, Clarke et al., 2007). Recently, the global chondrichthyan catch has been increasingly dominated by rays (Dulvy et al., 2014a). This increase is likely the result of (a) improved reporting, and (b) declines in shark catches due to stronger national and international regulations (Dulvy et al., 2014a). Rays, like most chondrichthyans, have intrinsically low biological productivity, due to their slow growth, late maturity, long generation times, and low fecundity, and are therefore slow to recovery from population declines (Fowler et al., 2002). Most ray species (excluding deep sea skates) are exposed to high levels of intense and expanding fishing pressure due to their strong association with soft bottom habitats in shallow (<100 m depth) tropical and temperate coastal waters (Last et al., 2016). Many species of rays play an important trophic role in soft sediment ecosystems as bioturbators (Kyne and Bennett, 2002, White et al., 2013, Flowers et al., 2020), and their coastal habitats are under threat from additional anthropogenic influences (Dulvy et al., 2016, Compagno and Cook, 1995).

Wedgefish (Family Rhinidae, 10 species), and giant guitarfish (Family Glaucostegidae, 6 species), are large (maximum size 300 cm total length, TL) benthic rays collectively referred to as shark-like rays (Last et al., 2016). They are found throughout the Indo-Pacific, Indian and Atlantic-Mediterranean ocean in shallow, coastal waters adjacent to coral reefs (Last et al., 2016). Shark-like rays are mainly caught as bycatch in fishing gears such as trawl nets, pelagic and bottom long lines, purse seine nets, and gillnets, and are typically retained as valuable by-products of opportunistic catch (Moore, 2017, Jabado, 2018). Shark-like ray fins are considered the highest grade in the international shark fin trade, which is likely the key driver for their retention in coastal fisheries (Keong, 1996). There are few documented targeted fisheries for these species (White and Dharmadi, 2007). Wedgefish and giant guitarfish are experiencing significant declines throughout their entire ranges (Kyne et al., 2019), and all but one species of wedgefish and giant guitarfish species are classified as Critically Endangered on the International Union for Conservation of Nature’s (IUCN) Global Red List of Threatened Species in 2019 (Kyne et al., 2020). These species were listed on Convention of International Trade of Endangered Species of Wild Fauna and Flora (CITES) Appendix II in 2019 (CoP18), which aims to ensure that the international trade of products from wedgefish and giant guitarfish come from sustainable sources (CITES, 2019).

Similar to the global decline of sawfishes and angel sharks (Lawson et al., 2020, Dulvy et al., 2016), depletion of wedgefish and giant guitarfish likely began many decades ago, driven by the incidental catch in fisheries and the high value of fins in the international trade (Clarke et al., 2006, Keong, 1996, Harrison and Dulvy, 2014). Quantifying the onset and extent of decline of these data-limited species is difficult, due to depletions occurring before independent scientific monitoring, poor fisheries and trade reporting resulting in little species-specific data, and lack of conservation awareness (Lawson et al., 2020, Dulvy et al., 2016). Wedgefish and giant guitarfish have been inferred to have a higher than average population productivity compared to other chondrichthyans, and therefore can potentially recover from population declines more rapidly than other threatened species (D’Alberto et al., 2019). However, achieving this will require significant reductions in fishing mortality (D’Alberto et al., 2019). Yet there is little information available on shark-like ray and stingray species’ historical and contemporary interactions with fisheries, which may hinder the development of management and enforcement efforts.

Indonesia is the world’s largest contemporary elasmobranch fishing nation, accounting for ca. 13% of reported global elasmobranch catch (Jaiteh et al., 2017, Blaber et al., 2009). It is also the third largest exporter of shark fins in regards to quantity, with an average of 1,235 tonne and sixth largest in value, worth an average of US$10 million per year (Dent and Clarke, 2015). Apart from the fin trade, elasmobranch meat is also an important source of protein for communities in South East Asia (Ahmad et al., 2016), and a large volume of elasmobranch products, particularly stingrays, continue to be exported to regional markets such as Singapore (Clark-Shen et al., 2021). In Indonesia, wedgefishes and guitarfishes are caught as bycatch in a variety of fisheries, but they are also specifically targeted in the tangle net fishery, with *Rhynchobatus australiae* the main target species. The fishery also lands stingrays and sharks as opportunistic catch (Keong, 1996, White and Dharmadi, 2007). This fishery uses large mesh (50 – 60 cm) bottom-set nylon gillnets, to capture large elasmobranchs by entanglement, on sandy or muddy substrates between 25 – 45 m depth (Amir, 1988). Fishing vessels in this fishery are typically refurbished 10 – 22 meter (30 – 40 tonnes) long wooden purse seine vessels (Amir, 1988). They have an approximate net load capacity of 15 tonnes, and are equipped with a diesel powered inboard engine and a small freezer to hold fins (Amir, 1988). In Indonesia, this fishery is referred to as “jaring liongbun” [= guitarfish gillnet] and “jaring cucut” [= shark gillnet].

The first records of this fishery are from Aru Island in the mid-1970s, from which the fishery rapidly expanded throughout Indonesian waters (Keong, 1996). It reached peak fishing capacity in 1987 with 500 active vessels and began operating at other ports, before declining by 80% over a ten year period (Amir, 1988, Dharmadi and Kasim, 2010). Declines of *Rhynchobatus* spp. began around 1992 according to local fishers that operate bottom set gillnets in the Aru-Arafura Sea (Jaiteh et al., 2016). The reductions in vessels operating in the tangle net fishery suggests that populations of the target species had declined and made the fishery economically unviable. The high value of wedgefish and giant guitarfish fins was a particularly strong driver for this fishery, with the fins worth approximately 1.5 times more than those from other species (Keong, 1996). More recently, high valued leather products from stingrays, which appear to be increasing in demand (Karthikeyan et al., 2009, Sahubawa et al., 2018), drives the retention of large stingrays. There is strong anecdotal evidence of declines of wedgefish and giant guitarfish in some areas of Indonesia as a result of this fishery (Amir, 1988, Keong, 1996). However, there are no catch and size composition data available for this fishery, and the fishery as a whole is poorly defined and little understood.

To achieve sustainable use of these species, managers and conservation practitioners need to understand their population status, risk exposure, and resilience to fishing pressure and other threats. This requires data on fisheries catch composition, changes in relative abundance, and their interactions with fisheries (Simpfendorfer et al., 2011, Jabado, 2018). The main aims of this paper were to (1) examine the species, size, and sex composition of the landed catch of Indonesian tangle net fishery in 2001 – 2005, and document changes over time in abundance and species compositions, (2) and to examine the potential consequences of these trends for future fisheries management and species conservation. Information of the Indonesian tangle net fishery can be used to inform the basis for the development of local and international management plans and conservation action for threatened rays.

## 2. Materials and Methods

### 2.1. Muara Angke landing port surveys

Between April 2001 and December 2005, elasmobranch landed catches from the tangle net fishery were recorded at the Muara Angke landing port (North Jakarta, Indonesia) (Fig. 1) and the adjacent village, where post-landing processing of fish occurred. Landing port surveys were conducted on 18 occasions, and for each visit the landing port was surveyed for 1 – 4 consecutive days, resulting in a total 53 sampling days (see Supplementary Table S1). Meanwhile, data on the nature of the products retained, their use, value, and export destination were collected by fisheries officers from the Ministry of Marine Affairs and Fisheries (MMAF). The prices reported for these products have been converted to Indonesian Rupiah (IDR) in 2021 values to account for inflation (www.inflationtool.com; 1 IDR 2004 = 2.54 IDR 2021; 1 IDR 2005 = 2.38 IDR 2021), and to US Dollar (USD) prices using an online currency converter (www.xe.com/currencyconverter/; $1 USD = 13,957.10 IDR as of January 2021).

**Figure 1.**
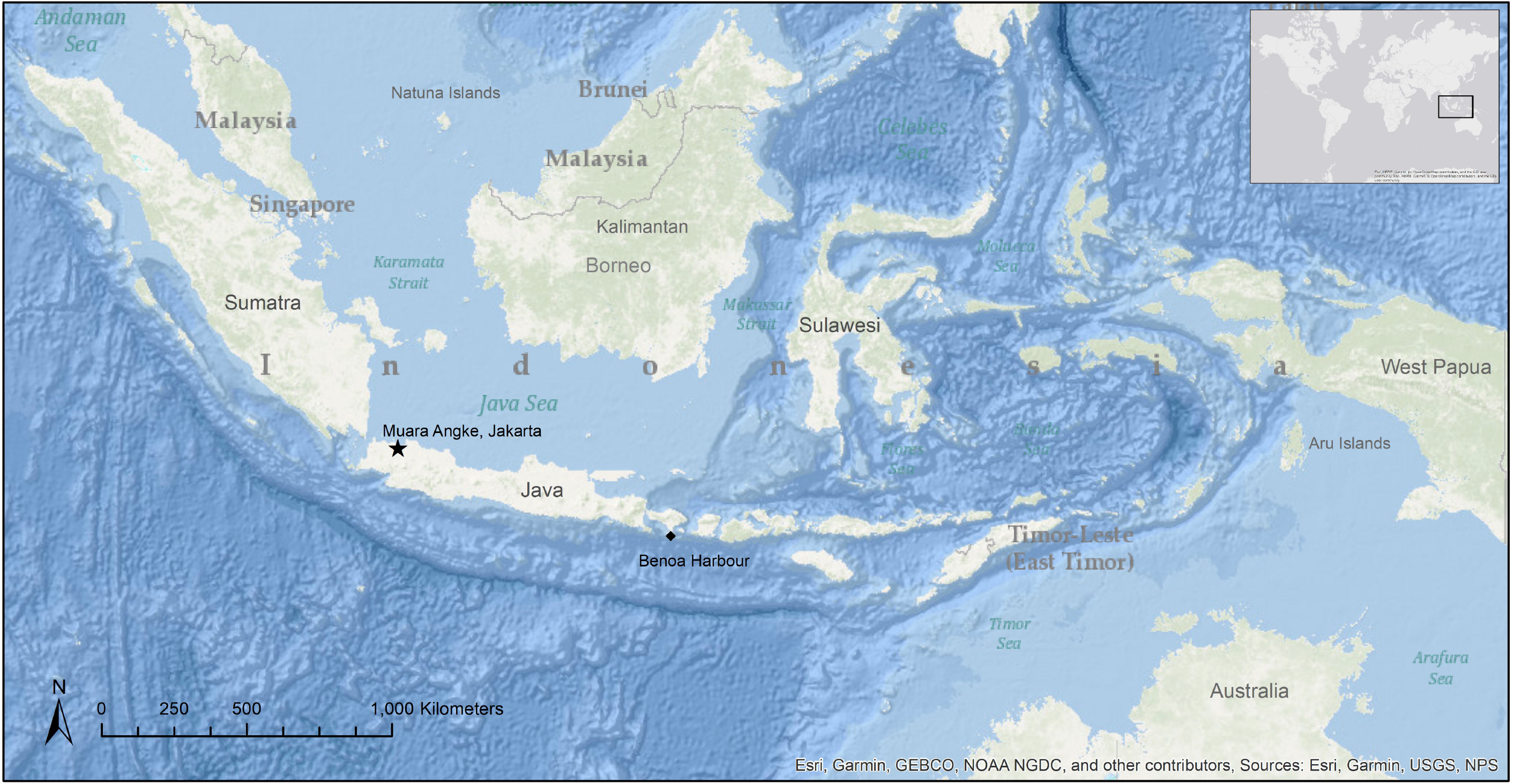
The location of the Muara Angke landing site and processing village in Jakarta (star), and Benoa Harbour on Bali (triangle), where the frozen shipment of landed catch was originally landed before relocated to processing village near Muara Angke, Indonesia. The map was created using ArcGIS software by Esri (Version 10.4.1; www.esri.com). Map Sources: Esri, USGS, NOAA, Garmin, NPS.

The number of each species landed from a tangle net vessel was recorded. Due to the large number of landings and time constraints on each day surveyed, the number of specimens, biological data and measurements could not be taken from all elasmobranches present. Only specimens that could be accessed were surveyed, so randomised selection for sex/size was not possible. At the Muara Angke landing port, catch composition could only be recorded for a brief period while the vessels were being unloaded (Fig. 2a, b). After the catches were unloaded, elasmobranchs were taken to the adjacent village processing area, located less than a kilometre from the fishing port. Here the large elasmobranchs from the tangle net fishery were typically taken to one of the four processing ‘houses’ (Fig. 2d). Similar data could be obtained at the village processing area, often from the previous day’s landings, but it was not possible to determine how many vessels they originated from if more than one vessel had landed in the previous two days. Species and size composition data was more readily collected during the unloading from the vessel at the fishing port. On days when catches were recorded in Muara Angke landing port, catches were not examined again in the village processing area. Due to the relatively low number of landings observed per trip, this issue was rarely encountered (Table S1). On one occasion, the landings from tangle net fishing vessels operating in the Banda and Arafura Seas that land at Benoa Harbour, Bali (Fig. 1), were observed and recorded in the Muara Angke processing village. These catches arrived into the village processing area by freezer truck direct from Bali.

**Figure 2.**
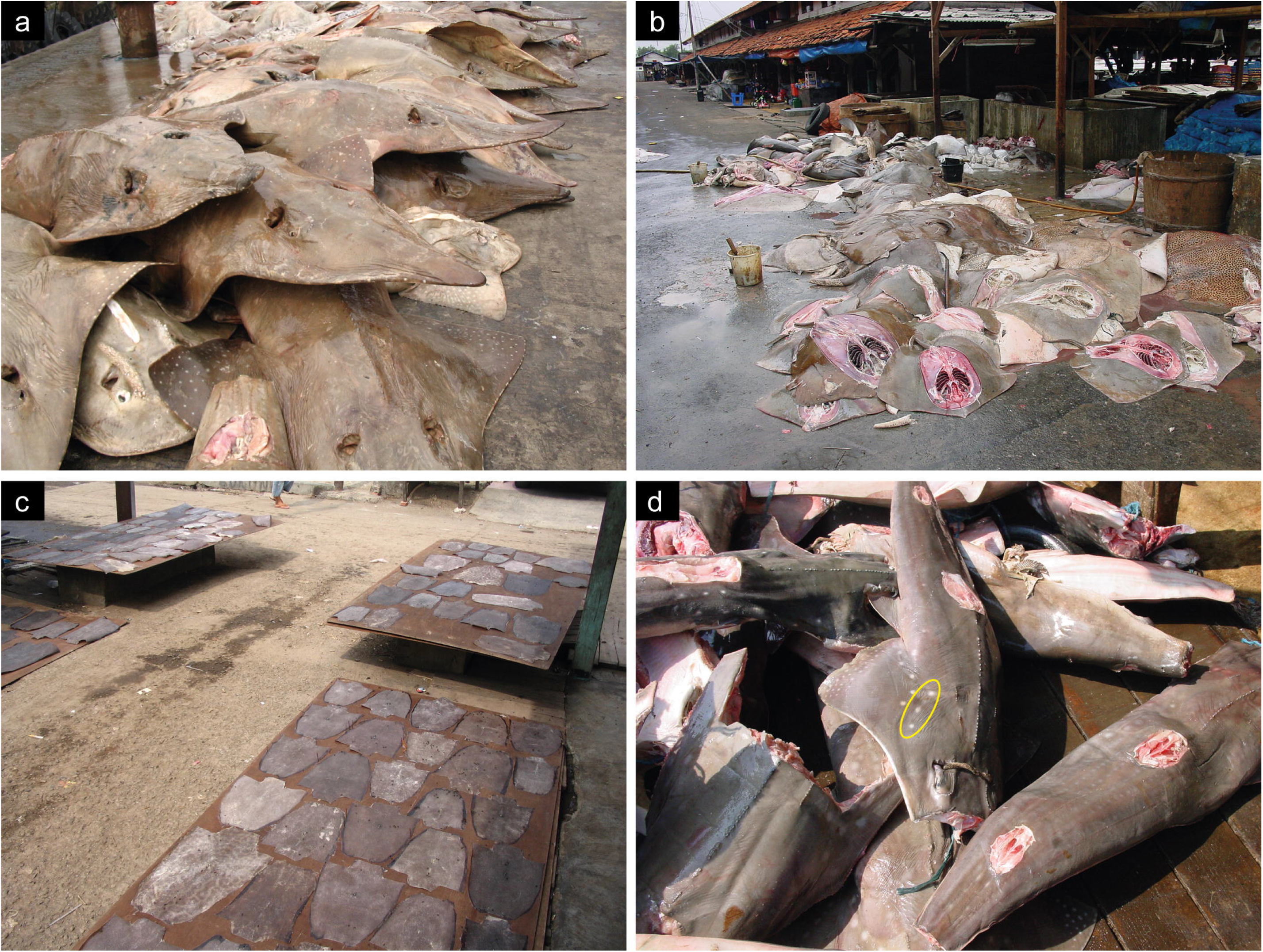
Tangle net fishery catches at Muara Angke landing site, Jakarta Indonesia: (a) Large bottlenose wedgefish *Rhynchobatus australiae* unloaded from tangle net vessels at the port; (b) Large stingrays being processed at the adjacent village processing area; (c) Drying ray skins which will be used to make stingray leather products such as wallets and belts; (d) Wedgefish landings from Arafura Sea at the village processing area – *Rhynchobatus australiae* in centre of image highlighting the line of three white spots (yellow circle) diagnostic in this species. Photo credits: W.T.W.

Often individual tangle net vessels would come into Muara Angke port once a month, and on three occasions it was possible to document the entire landed catch from three tangle net vessels. The entire landed catch from three vessels were recorded in Muara Angke landing site in July 2004, October 2004 and two in October 2005, and referred to as MA-SKR-170704, PV-PK-061004, and PV-UK-051005, respectively (Table 1). The landed catches from the vessels PV-KA-051005 and PV-UK-051005 were recorded on the same day. However, not all catch was able to be examined for PV-UK-051005, and the recorded catches for PV-KA-051005 and PV-UK-051005 were combined and assigned the vessel identifier code of PV-CC-051005 (Table 1).

**Table 1.**
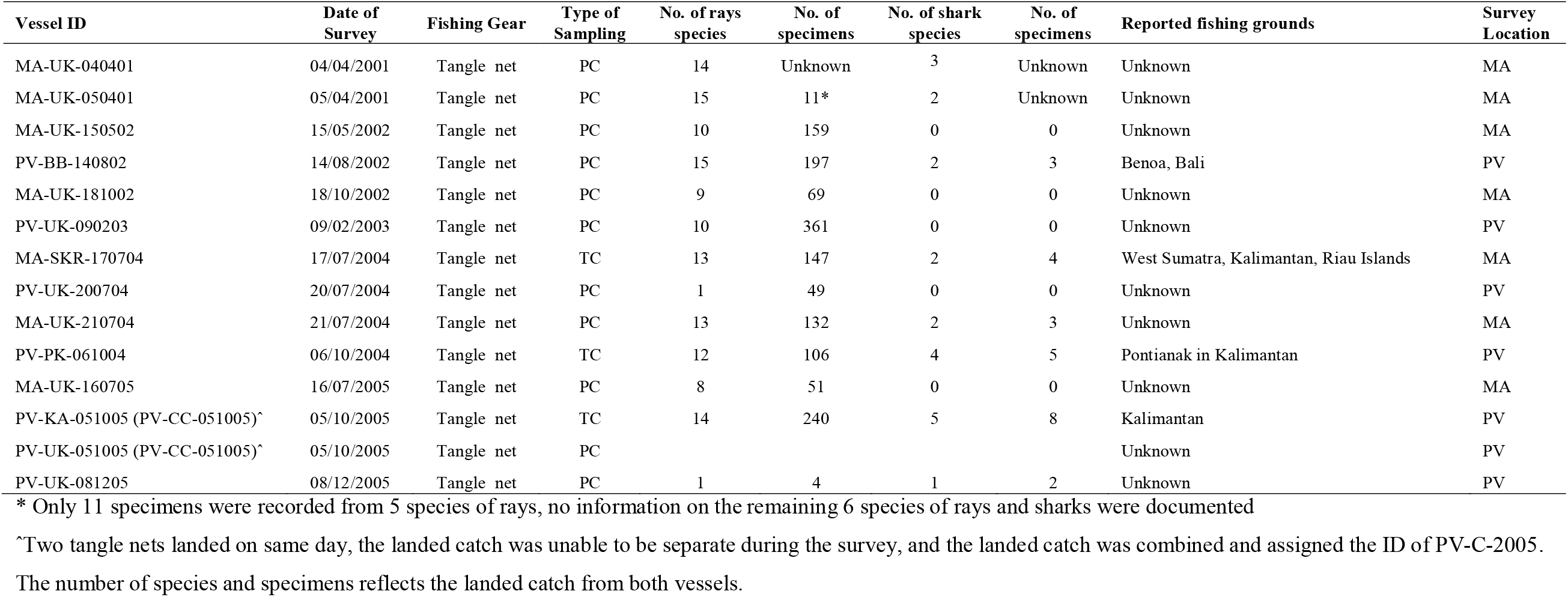
Summary table of the recorded tangle net vessels landed in Muara Angke landing site, Jakarta Indonesia in April 2001 – December 2005, including a vessel identifier (Vessel ID), date of survey, type of fishing gear, type of sampling (partial landed catch, PC; total landed catch, TC), number (No.) of ray and shark species and specimens surveyed, the reported fishing grounds, and survey location (Muara Angke landing site, MA; the nearby processing village, PV).

| Vessel ID | Date of Survey | Fishing Gear | Type of Sampling | No. of rays species | No. of specimens | No. of shark species | No. of specimens | Reported fishing grounds | Survey Location |
| --- | --- | --- | --- | --- | --- | --- | --- | --- | --- |
| MA-UK-040401 | 04/04/2001 | Tangle net | PC | 14 | Unknown | 3 | Unknown | Unknown | MA |
| MA-UK-050401 | 05/04/2001 | Tangle net | PC | 15 | 11* | 2 | Unknown | Unknown | MA |
| MA-UK-150502 | 15/05/2002 | Tangle net | PC | 10 | 159 | 0 | 0 | Unknown | MA |
| PV-BB-140802 | 14/08/2002 | Tangle net | PC | 15 | 197 | 2 | 3 | Benoa, Bali | PV |
| MA-UK-181002 | 18/10/2002 | Tangle net | PC | 9 | 69 | 0 | 0 | Unknown | MA |
| PV-UK-090203 | 09/02/2003 | Tangle net | PC | 10 | 361 | 0 | 0 | Unknown | PV |
| MA-SKR-170704 | 17/07/2004 | Tangle net | TC | 13 | 147 | 2 | 4 | West Sumatra, Kalimantan, Riau Islands | MA |
| PV-UK-200704 | 20/07/2004 | Tangle net | PC | 1 | 49 | 0 | 0 | Unknown | PV |
| MA-UK-210704 | 21/07/2004 | Tangle net | PC | 13 | 132 | 2 | 3 | Unknown | MA |
| PV-PK-061004 | 06/10/2004 | Tangle net | TC | 12 | 106 | 4 | 5 | Pontianak in Kalimantan | PV |
| MA-UK-160705 | 16/07/2005 | Tangle net | PC | 8 | 51 | 0 | 0 | Unknown | MA |
| PV-KA-051005 (PV-CC-051005)^ | 05/10/2005 | Tangle net | TC | 14 | 240 | 5 | 8 | Kalimantan | PV |
| PV-UK-051005 (PV-CC-051005)^ | 05/10/2005 | Tangle net | PC |  |  |  |  | Unknown | PV |
| PV-UK-081205 | 08/12/2005 | Tangle net | PC | 1 | 4 | 1 | 2 | Unknown | PV |

\* Only 11 specimens were recorded from 5 species of rays, no information on the remaining 6 species of rays and sharks were documented
^Two tangle nets landed on same day, the landed catch was unable to be separate during the survey, and the landed catch was combined and assigned the ID of PV-C-2005.
The number of species and specimens reflects the landed catch from both vessels.

Following the methods described above, catch composition of elasmobranchs from other fisheries were also recorded during the Muara Angke landing site surveys. This included landings from small-mesh gillnet (<20 cm mesh size) fisheries, Java Sea and Arafura Sea trawl fisheries, southern Java trammel net fishery, and various hand- and long-line fisheries, which were operating out of the landing ports surveyed [see White and Dharmadi (2007)]. This allowed the comparison of the size composition of species between the tangle net fishery and the other fisheries interacting with the same species. Similar to the tangle net fishery landings, only landed catch that could be accessed when a vessel was unloading was able to be surveyed, and randomised selection was not possible.

### 2.2. Biological data

When possible, the disc width (DW) for the Dasyatidae, Myliobatidae, Aetobatidae, Gymnuridae and Rhinopteridae, and total length (from the tip of the snout to the tip of the upper lobe of the caudal fin; TL) for the sharks and shark-like rays (Pristidae, Glaucostegidae and Rhinidae) were measured to the nearest 1 cm, and sex recorded. As the shark-like rays were typically landed without fins, an estimated TL was recorded when animals were landed without fins. Occasionally, the removal of fins from these rays occurred following landing, and after weighing of specimens. Total weight (TW) of whole individuals (fins attached and not gutted) was recorded to the nearest g or kg (depending on the size of the individual), however, the vast majority of rays and sharks could not be weighed at the landing port. When large numbers of similar sized individuals were observed, measurements were taken from a subset of whole individuals that could be accessed, and used to estimate DW, TL and TW for the remaining unmeasured individuals. For specimens that were not weighed but had length measurements, the weight of individual specimens were calculated using the length to weight conversion equations (see Supplementary Table S2). For species where a length-weight equation was not available, the estimated weight was calculated using a conversion equation from a morphologically-similar species (Table S2). In instances when the size of individuals for a particular species was not recorded, the weight was estimated using the average weight of the individuals for that species. Total weight was then determined for each species landed in the fishery. Details on the reproductive biology of each species recorded were previously reported in White and Dharmadi (2007).

### 2.3. Species identification

Species were identified using the keys in Carpenter and Niem (1998) and Last and Stevens (1994), with taxonomic nomenclature updated using Last et al. (2016) and Last and Stevens (2009). The identity of a subsample of *Rhynchobatus* species caught in the tangle net fishery was also further verified by genetic analysis (Giles et al., 2016), and from images that used recently determined colour pattern differences between species (Last et al., 2016). The key colour pattern difference used to differentiate *R. australiae* from its closest regional congeners was the pattern of white spots around the dark pectoral spot present in all but the largest individuals (Fig. 2d). In *R. australiae*, there is a line of 3 white spots located adjacent to the black pectoral spot, or its usual position if faded (Last et al., 2016). In two large pregnant females which possessed no white spots or black pectoral spots, due mainly to their poor condition, the typical *R. australiae* spot pattern was evident in late-term embryos allowing for confirmation of their identity. *Maculabatis macrura* was only recently recognised as valid and distinct from *Maculabatis gerrardi* (Last et al., 2016), therefore our data could not be retrospectively confirmed as either or both species (Last et al., 2016). Although these records could constitute either species, herein we refer to these species as the prevailing name used for the whitespotted ray species in Indonesia, *M. gerrardi*.

### 2.4. Data analysis

Species composition of each landing was expressed as percentage of the total number of individuals by both the number and weight for each species recorded at the landing port and processing village. Minimum, maximum and mean ± standard error (S.E.) for DW, TL and TW are reported for each species. Size frequency histograms for the most abundant species were produced.

### 2.5. Ethics Statement

All marine life examined in this study were landed from fisheries in Indonesia and were already dead upon inspection. Permission to undertake surveys in Indonesia was granted by the Research Centre for Fisheries Management and Conservation in Jakarta as part of collaborative projects funded by the Australian Centre for International Agricultural Research (ACIAR project codes FIS/2000/062 and FIS/2003/037).

## 3. Results

### 3.1. Species and size composition of the fishery

Across 18 sampling trips, totalling 53 survey days from April 2001 – December 2005, tangle net vessel landings were recorded in Muara Angke landing site eight times, and within the village processing area seven times, including one frozen shipment from Benoa Harbour, Bali (Table 1; Table S1). From discussions with a fleet manager, thirteen vessels were reported to be operational in the tangle net industry in 2004, fishing in waters around Borneo, Sulawesi and as far as West Papua (Fig 1). Only vessels that landed at Muara Angke landing site or processing village during the 2001-2005 surveys were sampled (Table 1).

A total of 1,559 elasmobranchs were recorded from tangle net fishery landings at Muara Angke landing port. This comprised 1,526 batoids (98.3% of the catch), of 24 species from seven families (Table 2). The most abundant family was Dasyatidae, contributing 72.5% to the total number of elasmobranchs recorded, followed by the family Rhinidae, and comprised 20.8% of total observed catch. Only 25 sharks were recorded, with nine shark species from four families (Table 2). Combining the total landed catch of the four Indonesian tangle net vessels, of which the catch was fully documented from Muara Angke landing site, the most abundant species was *M. gerrardi*, followed by *R. australiae, P. ater, R. ancylostoma, A. ocellatus, H. uarnak*, and *P. jenkinsii* (Fig. 5). In total, eighteen other species of elasmobranchs, comprising 10 ray species and 8 shark species, were also recorded but in low numbers. As not all individuals could be counted when landed, and in many cases the numbers were estimated, the numbers presented represent an underestimation of the total number of individuals caught during the survey period.

**Table 2.** Species composition, number of individuals of a species observed (no.) and overall percentage (% by no.) of the catch for the elasmobranchs caught by the tangle net fishery, and landed in Muara Angke landing site, Jakarta Indonesia in April 2001–December 2005. The number of recorded specimens (No.), percentage of total specimens (% by No.), observed minimum (Min. size), observed maximum size (Max. size), observed mean (± S.E.) size (DW/TL cm), minimum (Min. weight) and maximum (Max. weight) estimated total weight (kg) are reported for each species, with the International Union for Conservation of Nature’s (IUCN) Red List of Threatened Species status (as of August 2020). IUCN categories are CR, Critically Endangered; EN, Endangered; VU, Vulnerable; NT, Near Threatened; LC, Least Concern; DD, Data Deficient. Dashed lines indicate species presence was recorded in landings, but data was not able to be documented.

| Family | Scientific name | Common name | IUCN Listing | Year Assessed | No. | % by No. | Min. size | Max. size | Mean Size | ± S.E. | Min. weight | Max. weight | Mean Weight | ± S.E. |
| --- | --- | --- | --- | --- | --- | --- | --- | --- | --- | --- | --- | --- | --- | --- |
| Pristidae | *Pristis pristis* | Large-tooth sawfish | CR | 2013 | 2 | 0.128 | -- | 420.0 | -- | -- | -- | 220.3 | -- | -- |
| Glaucostegidae | *Glaucostegus thouin* | Clubnose guitarfish | CR | 2019 | -- | -- | -- | -- | -- | -- | -- | -- | -- | -- |
|  | *Glaucostegus typus* | Giant guitarfish | CR | 2019 | 14 | 0.898 | 170.0 | 260.0 | 206.0 | 13.15 | 14.86 | 51.36 | 27.57 | 5.543 |
| Rhinidae | *Rhina ancylostoma* | Bowmouth guitarfish | CR | 2019 | 57 | 3.656 | 130.1 | 270.0 | 190.2 | 22.97 | 18.68 | 168.4 | 77.38 | 24.29 |
|  | *Rhynchobatus australiae* | Bottlenose wedgefish | CR | 2019 | 238 | 15.27 | 190.0 | 300.0 | 282.1 | 4.662 | 30.21 | 118.7 | 100.9 | 4.181 |
|  | *Rhynchobatus palpebratus* | Eye-brow wedgefish | NT | 2019 | 30 | 1.924 | -- | -- | -- | -- | -- | -- | -- | -- |
| Dasyatidae | *Bathytoshia lata* | Brown stingray | LC | 2007 | 1 | 0.064 | -- | 202.0 | -- | -- | -- | 300.0 | -- | -- |
|  | *Himantura leoparda* | Leopard whiplay | VU | 2015 | 31 | 1.988 | 83.20 | 120.0 | 98.72 | 3.016 | 14.37 | 39.45 | 23.61 | 2.061 |
|  | *Himantura uarnak* | Coach whiplay | VU | 2015 | 57 | 3.656 | 42.60 | 147.6 | 99.50 | 6.204 | 4.015 | 69.83 | 28.57 | 4.189 |
|  | *Himantura undulata* | Honeycomb whiplay | VU | 2011 | 1 | 0.064 | -- | 112.8 | 112.8 | -- | 33.27 | -- | 33.27 | -- |
|  | *Maculabatis astra* | Blackspotted whiplay | LC | 2015 | 4 | 0.257 | -- | 79.00 | 79.00 | -- | 13.26 | -- | 13.26 | -- |
|  | *Maculabatis gerrardi* | Whitespotted whiplay | VU | 2004 | 194 | 12.44 | 62.70 | 89.50 | 75.85 | 1.036 | 6.560 | 19.40 | 12.01 | 2.410 |
|  | *Megatrygon microps* | Smalleye stingray | DD | 2015 | 1 | 0.064 | -- | 174.8 | -- | -- | -- | 111.3 | -- | -- |
|  | *Pastinachus ater* | Broad cowtail ray | LC | 2015 | 199 | 12.76 | 86.00 | 149.0 | 114.3 | 2.618 | 15.74 | 71.67 | 37.98 | 2.332 |
|  | *Pateobatis fai* | Pink whiplay | VU | 2015 | 264 | 16.93 | 70.50 | 168.4 | 110.9 | 4.595 | 9.101 | 100.4 | 36.45 | 4.143 |
|  | *Pateobatis jenkinsii* | Jenkin's whiplay | VU | 2015 | 187 | 11.99 | 59.20 | 138.4 | 82.59 | 1.791 | 5.621 | 58.47 | 14.78 | 1.207 |
|  | *Pateobatis uarnacoides* | Whitenose whiplay | VU | 2004 | 125 | 8.018 | 51.70 | 118.8 | 91.23 | 4.206 | 3.869 | 38.38 | 20.42 | 2.390 |
|  | *Taeniurops meyeri* | Blotched stingray | VU | 2015 | 51 | 3.271 | 62.80 | 164.0 | 116.8 | 5.332 | 6.615 | 93.37 | 40.25 | 4.713 |
|  | *Urogymnus asperrimus* | Porcupine ray | VU | 2015 | 5 | 0.321 | 76.50 | 103.4 | 89.95 | 9.511 | 11.40 | 26.17 | 18.78 | 5.222 |
|  | *Urogymnus granulatus* | Mangrove whiplay | VU | 2015 | 10 | 0.641 | 97.20 | 141.0 | 118.8 | 6.300 | 22.07 | 61.55 | 39.72 | 5.728 |
| Gymnuridae | *Gymnura zonura* | Zonetail butterfly ray | VU | 2006 | 7 | 0.449 | 70.50 | 91.60 | 79.78 | 3.036 | 2.442 | 5.466 | 3.657 | 0.4433 |
| Aetobatidae | *Aetobatus ocellatus* | Spotted eagle ray | VU | 2015 | 45 | 2.886 | 108.9 | 214.4 | 138.6 | 3.994 | 19.37 | 135.4 | 41.43 | 4.151 |
| Myliobatidae | *Aetomylaeus vespertilio* | Ornate eagle ray | EN | 2015 | 11 | 0.706 | 146.2 | 240.0 | 187.5 | 17.157 | 45.11 | 187.1 | 100.6 | 26.66 |
| Carcharhinidae | *Carcharhinus amboinensis* | Pigeye shark | DD | 2005 | -- | -- | -- | -- | -- | -- | -- | -- | -- | -- |
|  | *Carcharhinus obscurus* | Dusky shark | VU | 2007 | 2 | 0.128 | -- | -- | -- | -- | -- | -- | -- | -- |
|  | *Carcharhinus leucas* | Bull shark | NT | 2005 | 2 | 0.128 | -- | -- | -- | -- | -- | -- | -- | -- |
|  | *Carcharhinus limbatus* | Common blacktip shark | NT | 2005 | 6 | 0.385 | -- | -- | -- | -- | -- | -- | -- | -- |
|  | *Galeocerdo cuvier* | Tiger shark | NT | 2005 | 4 | 0.257 | -- | -- | -- | -- | -- | -- | -- | -- |
| Ginglymostomatidae | *Nebrius ferrugineus* | Tawny nurse shark | VU | 2003 | 3 | 0.192 | -- | -- | -- | -- | -- | -- | -- | -- |
| Hemigaleidae | *Hemipristis elongata* | Fossil shark | VU | 2015 | 3 | 0.192 | 109.6 | 122.9 | 116.3 | 4.702 | 5.719 | 25.14 | 6.989 | 0.8984 |
| Sphyrnidae | *Sphyrna lewini* | Scalloped hammerhead | CR | 2019 | 3 | 0.192 | -- | 175.4 | 175.4 | -- | -- | -- | 25.14 | -- |
|  | *Sphyrna mokarran* | Great hammerhead | CR | 2019 | 2 | 0.128 | -- | -- | -- | -- | -- | -- | -- | -- |

#### 3.1.1. Family Pristidae

Two largetooth sawfish *Pristis pristis* were recorded from the Benoa Harbour, Bali landings in August 2002. It is likely the sawfish were caught in the Arafura or Banda Sea region (Fig. 1). Both individuals were adult males and ca. 420 cm TL and had an estimated total landed weight of 220 kg (Table 2).

#### 3.1.2. Family Glaucostegidae

Two species of giant guitarfish were recorded in Muara Angke landing site: the clubnose guitarfish *Glaucostegus thouin*, and giant guitarfish *Glaucostegus typus*. The presence of *G. thouin* in the fishery was recorded once in April 2001. However due to the logistics of accessing these rays upon unloading from the vessel and the decayed state of some specimens, the estimates of numbers or size were not possible for *G. thouin*, and it was unclear if it was a regular catch in the fishery. Fourteen *G. typus* were recorded on three occasions (Table 2). Of the subset of *G. typus* measured, seven were females, six males and one not sexed, with an estimated total landed weight of 386 kg (Table 2).

#### 3.1.3. Family Rhinidae

Three species of wedgefish were recorded: the bowmouth guitarfish *Rhina ancylostoma*, the eyebrow wedgefish *Rhynchobatus palpebratus*, and *R. australiae. Rhina ancylostoma* were recorded on seven occasions over the sampling period (Table S3). The landed catch of *R. ancylostoma* comprised 15 females, 10 males, and 32 specimens counted but not sexed, with an estimated total landed weight of 4.4 tonnes (Table 2). Females ranged from 139 – 250 cm TL and 22.9 – 133.6 kg, and males ranged from 130 – 260 cm TL and 18.7 – 150.3 kg. One unsexed specimen was measured at 270 cm TL and 168 kg. The main target species of the tangle net fishery, *R. australiae* comprised the largest component of wedgefishes and the second most abundant species recorded (Table 2; Fig. 5). On one occasion, approximately 7.1 tonne of *R. australiae* was landed from a single tangle net vessel. *Rhynchobatus australiae* was recorded on eight occasions (Table S3) and a total of 238 individuals with an estimated total landed weight of 24 tonnes, comprising 99 females, 18 males and 121 unsexed individuals (Table 2). A subset of 29 individuals were measured, the majority of which were females and approximately 300 cm TL (Fig. 2a). Of the subset, 16 individuals of *R. australiae* examined internally were pregnant as reported in White and Dharmadi (2007).

A total of 16 *Rhynchobatus* spp. landed in the tangle net fishery had their identifications confirmed by genetic analysis [see Giles et al. (2016)]. Of these 16, two were from landings at Muara Angke and the remaining 14 from the Benoa Harbour landings. The 14 Benoa Harbour individuals consisted of five *R. palpebratus* [reported as *R. palpebratus/Rhynchobatus cf laevis* in Giles et al. (2016)] and nine *R. australiae*. The ratio of *R. palpebratus* to *R. australiae* determined from the genetic analysis (45:6) was used to estimate the species composition of the 100 *Rhynchobatus* individuals recorded in the Benoa Harbour landings on the 14^th^ August 2002. *Rhynchobatus palpebratus* was recorded on one occasion, with a total of 30 individuals but not sexed, measured or weighed (Table 2).

#### 3.1.4. Family Dasyatidae

Stingrays were present in every tangle net catch landed in Muara Angke. A total of 1130 stingrays, with an estimated mass of 30.2 tonnes, were recorded comprised of 13 species from eight genera (Table 2). The majority of the specimens caught of each species were near or at a larger size than their known size at maturity (Fig. 3, 4; see Supplementary Table S4). The most abundant stingray species were *P. fai* (9.6 tonnes; Fig. 3f), broad cowtail ray *Pastinachus ater* (7.5 tonnes; Fig. 3e), *M. gerrardi* (2.3 tonnes; Fig. 3d; Fig. 5), Jenkin’s whipray *Pateobatis jenkinsii* (2.7 tonnes; Fig. 4a), and whitenose whipray *Pateobatis uarnacoides* (2.5 tonnes; Fig. 4b). Other species that were recorded were the brown stingray *Bathytoshia lata*, leopard whipray *Himantura leoparda* (Fig. 3b), coach whipray *Himantura uarnak* (Fig. 3c), honeycomb whipray *Himantura undulata*, blackspotted whipray *Maculabatis astra*, smalleye stingray *Megatrygon microps*, blotched stingray *Taeniurops meyeni* (Fig. 4c) and porcupine ray *Urogymnus asperrimus. Maculabatis astra* was only recorded from the single Benoa Harbour landing, this species is only present in the far eastern portion of Indonesia off West Papua and is allopatric from *M. gerrardi* (Last et al., 2016). Specimens of *H. leoparda, H. uarnak, P. fai, P. uarnacoides* and *T. meyeni* specimens were close to the known maximum size (Fig. 3, 4).

**Figure 3.**
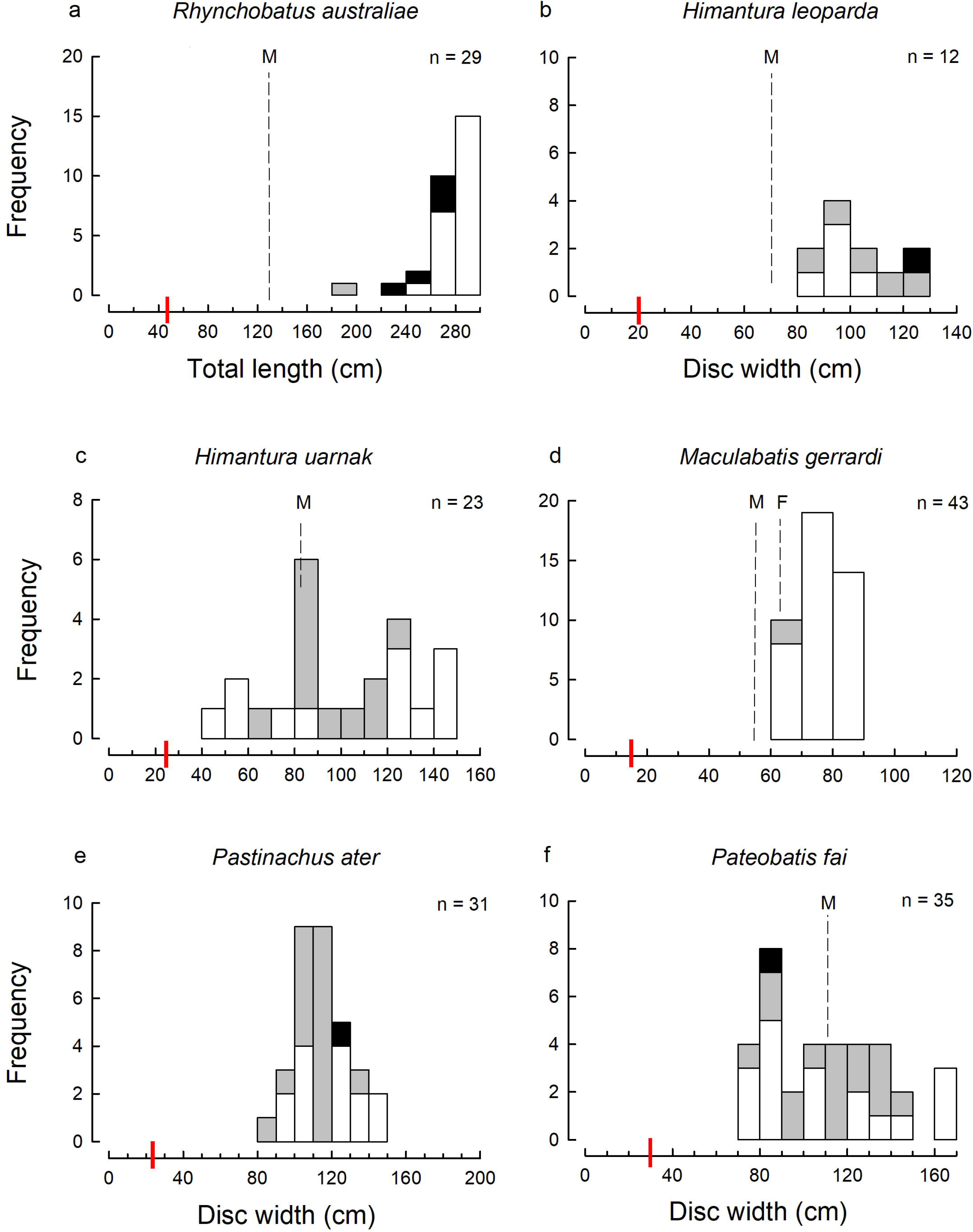
Size-frequency histograms of the most abundant ray species (represented by 10 or more measured individuals) in the tangle net landed catches from the Muara Angke landing site surveys 2001 – 2005: (a) *Rhynchobatus australiae*; (b) *Himantura leoparda*; (c) *Himantura uarnak*; (d) *Maculabatis gerrardi*; (e) *Pastinachus ater*; (f) *Pateobatis fai*. White bars denote females, grey bars males and black bars unsexed individuals; the total number (n) of individuals, size at birth (red solid line) and size at maturity (black dashed line; M, male; F, female) when known; the size scale bar (x-axis) extends to the maximum known size for each of the species. The species are placed in phylogenetic order from wedgefish through to eagle rays. Size at birth and size at maturity estimates are from Last et al. (2016).

**Figure 4.**
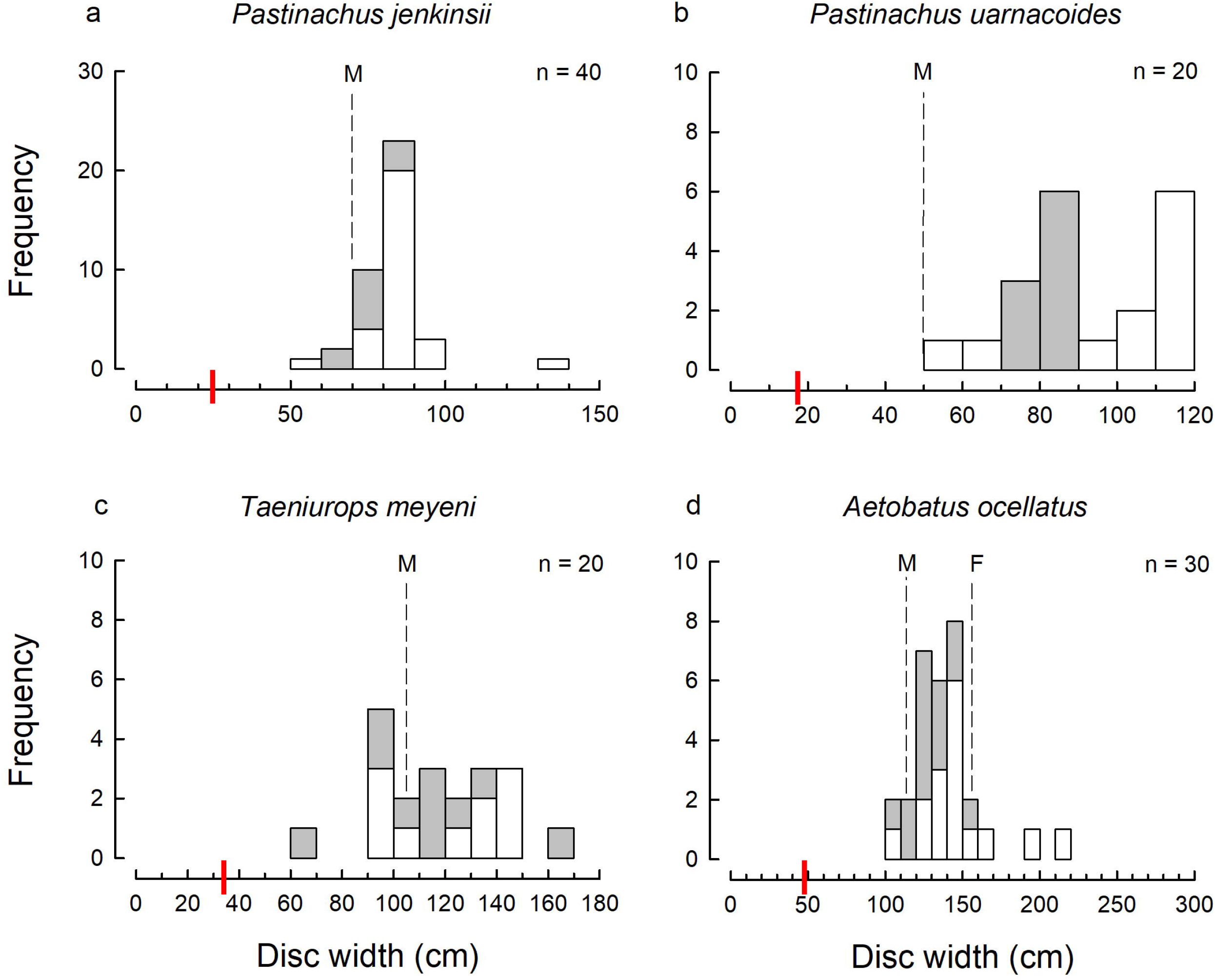
Size-frequency histograms of the most abundant ray species (represented by 10 or more measured individuals) in the tangle net catches from the Muara Angke landing site surveys 2001 – 2005: (a) *Pateobatis jenkinsii*; (b) *Pateobatis uarnacoides*; (c) *Taeniurops meyeni*; (d) *Aetobatus ocellatus*. White bars denote females, grey bars males and black bars unsexed individuals; the total number (n) of individuals, size at birth (red solid line) and size at maturity (black dashed line; M, male; F, female) when known; the size scale bar (x-axis) extends to the maximum known size for each of the species. The species are placed in phylogenetic order from wedgefish through to eagle rays. Size at birth and size at maturity estimates are from Last et al. (2016).

**Figure 5.**
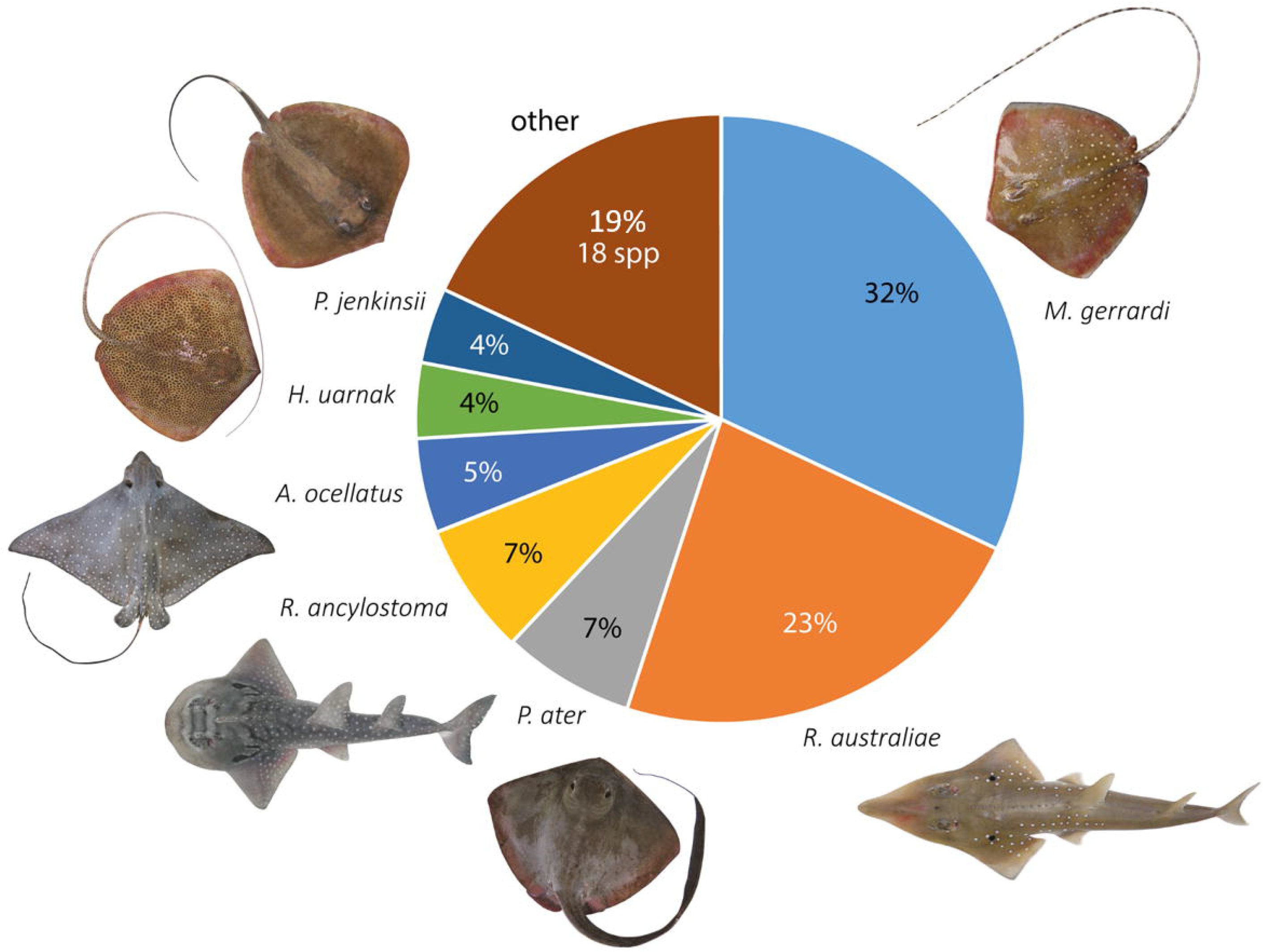
Overall species composition and percentage (%) of catch of the four Indonesian tangle net vessels (total n = 481), of which the landed catch was able to be fully documented from Muara Angke landing site, Jakarta Indonesia. The total landed catch from the four vessels consisted of *Maculabatis gerrardi* n = 155, *Rhynchobatus australiae* n = 112, *Rhina ancylostoma* n = 32, *Aetobatus ocellatus* n = 25, *Pateobatis jenkinsii* n = 18, *Himantura uarnak* n = 17, and for the other species n = 90. *MA*-SKR-170704 landed on 7^th^ July 2004; PV-PK-061004 landed on 16^th^ October 2004; PV-CC-051005 (combined landed catch of PV-KA-051005 and PV-UK-051005) landed on 5^th^ October 2005. Photo credit: W.T.W.

#### 3.1.5. Family Aetobatidae

One aetobatid species, the spotted eagle ray *Aetobatus ocellatus*, was recorded in the tangle net fishery on eight occasions (Table S3). A total of 45 individuals were observed, with an estimated total landed weight of 1.8 tonnes (Table 2). This comprised of 21 females, 22 males, and two unsexed specimens (Fig. 4d). *Aetobatus ocellatus* specimens were mainly caught close to or at a greater size than the known size at maturity (Fig. 4d; Table S4).

#### 3.1.6. Other families

Several other species from a range of families were recorded occasionally or in relatively small quantities. This included one species of Myliobatidae, the ornate eagle ray *Aetomylaeus vespertilio*, with an estimated landed catch of 1.1 tonnes, of which five were females, one male, and five unsexed individuals (Table 2). These specimens were all large, including one 160 kg female, and were recorded occasionally (Table S3), and comprised a small proportion of the total landed catch. A single Gymnuridae species was recorded, the zonetail butterfly ray *Gymnura zonura* with seven individuals recorded (Table 2). Sharks were a minor part of the catch and rarely observed. All of the shark species represented less than 1% of the total catch (Table 2). Of the small amount of sharks taken in the fishery, the species most commonly occurring in the catch were the common blacktip shark (*Carcharhinus limbatus*; n = 6), tiger shark (*Galeocerdo cuvier*; n = 4), and hammerheads (*Spyhrna* spp; n = 5) (Table 2; Table S3).

### 3.2. Products from the tangle net fishery

Data from MMAF indicated that the main products derived from the tangle net fishery catch were elasmobranch fins, leather made from ray skins, salted elasmobranch meat, and elasmobranch vertebrae. The most valuable product were fins from wedgefishes and giant guitarfishes. During the surveys in 2005, the quoted price for fins from sawfish, wedgefish and guitarfish were approximately 3 million IDR kg^-1^ wet weight (∼ 7,145,108 IDR / $512 USD per kg in 2021 terms). Any fins frozen on-board the tangle net vessels did not come through the Muara Angke landing site, but instead were directly exported through a different port to Hong Kong and Singapore.

Stingray skins, used to produce leather products, were the second most valuable product in the fishery (Fig. 2c). The species primarily used were from the genera *Himantura, Maculabatis, Pastinachus, Pateobatis* and *Urogymnus*, which together were a large component of the landed catch during the market surveys. *Pateobatis jenkinsii* was reported to be the most sought after stingray skin due to the row of enlarged thorns which extend down the midline of the body and tail. In 2005, the reported value of a 13 cm and 18 cm piece of stingray leather was 25,000 (= 59,543 IDR / $4.30 USD in 2021) and 35,000 IDR (= 83,360 IDR / $5.97 USD in 2021). Approximately 3000 – 4000 skins were estimated to be exported per month to the Philippines and Japan. Products from the stingray leather include wallets and belts, which were reported to be sold for approximately 290,000 IDR (= 690,694 IDR / $49.50 USD in 2021) to 500,000 IDR (= 1,190,851 IDR / $85.33 USD in 2021), as well as bags. Estimated prices for bags were not available.

Ray meat, wedgefish in particular, was considered to be good quality. As the catch was landed in a deteriorated condition, meat from the rays and sharks was salted and dried. In 2004, in the Muara Angke processing village, the buying price for wedgefish and guitarfish meat was 4,000 – 5,000 IDR kg^-1^ (= 10,143 – 12,679 IDR kg^-1^/ $0.73 – 0.91 USD kg^-1^ in 2021). For stingrays, the meat was valued between 2,000 – 3,500 IDR kg^-1^ (= 5,072 – 8,875 IDR kg^-1^/ $0.36 – 0.64 USD kg^-1^ in 2021). Salted meat was reported to be transported to West Java (Bandung, Bogor, Garut, Cianjur) and Central Java. Salted meat from wedgefish was reported to be sold for 10,000 – 12,000 IDR kg^-1^ (= 25,358 – 30,430 IDR kg^-1^/ $1.82 – 2.20 USD kg^-1^ in 2021), and for stingray meat 6,000 – 8,000 IDR kg^-1^ (= 15,215 – 20,287 IDR kg^-1^/ $1.09 – 1.45 USD kg^-1^ in 2021).

The cartilage, such as vertebrae, comprised a small part of the products from this fishery. In 2004, dried vertebrae were worth approximately 20,000 IDR kg^-1^ (= 50,717 IDR kg^-1^/ $3.63 USD kg^-1^ in 2021), which were then processed in Jakarta and exported through an undisclosed port to Korea and Japan.

### 3.3. Comparison of the size range of species from tangle net fishery and other fisheries

The gill nets used in the tangle net fishery captured wedgefish and guitarfish over 130 cm TL, and stingrays over 50 cm DW (Fig. 3), as a result of the large mesh size used (50 – 60 cm). The smallest recorded individual caught in this fishery was a *P. uarnacoides* of 51.7 cm DW and the largest recorded individual was a male sawfish, estimated to be 420 cm TL. Meanwhile, smaller size classes (<50 cm DW) for many ray species were captured in a range of fishing gears used in other fisheries operating in Indonesian waters during the 2001 – 2005 surveys, including trawl nets, hand- and long-lines, smaller mesh gillnets, and trammel nets (Fig. 6). These other fisheries greatly increased fishing selectivity for some species. For example, all size classes of *R. australiae* were captured; neonates (∼45 cm TL) were caught as bycatch in small mesh gillnets (Fig. 6a) while sub-adults (∼90 – 130 cm TL) were captured in the Java Sea trawl fishery, and the larger mature individuals (>170 cm TL) were taken in hand- and long-line fisheries, likely as by-products (Fig. 6a).

**Figure 6.**
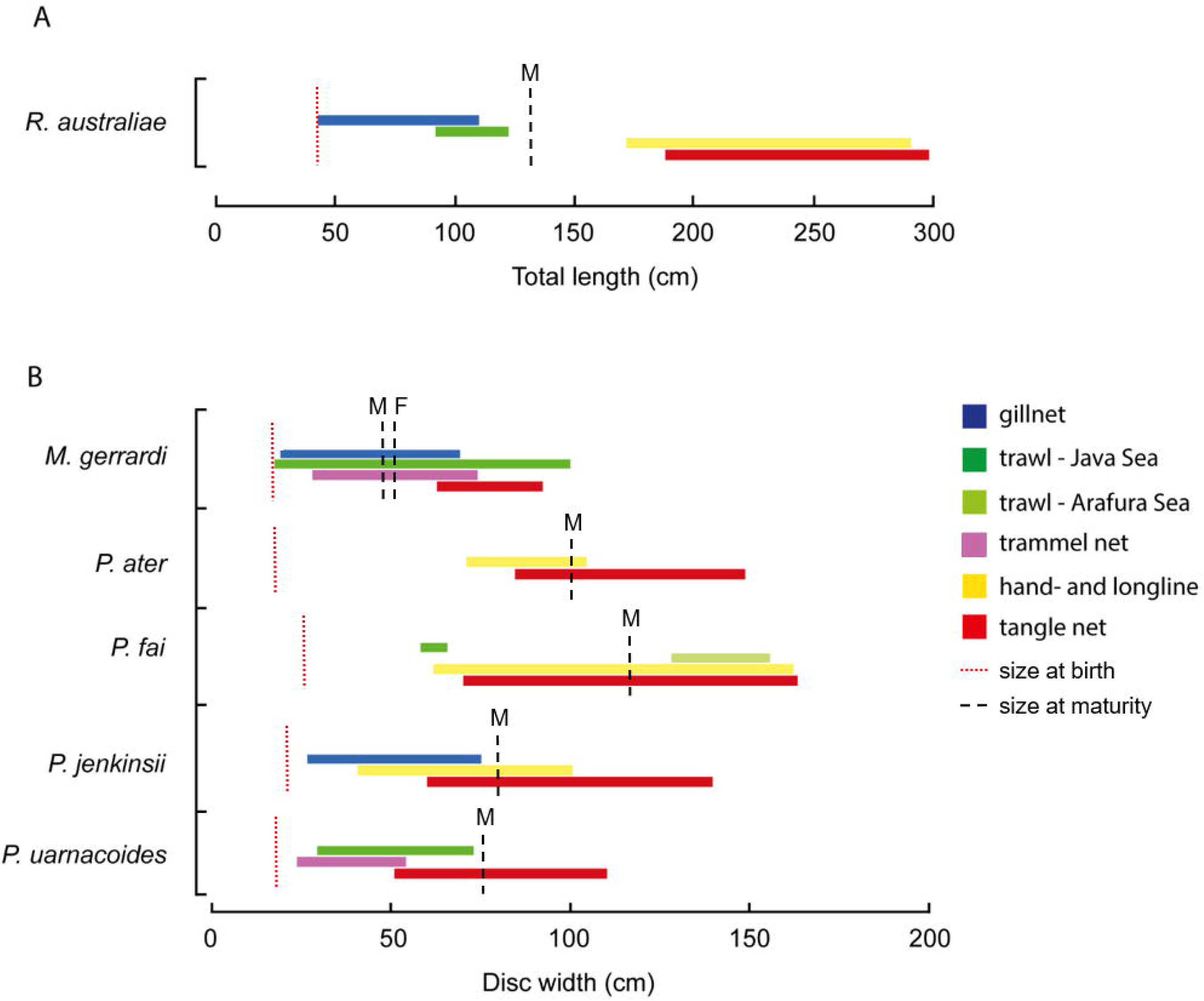
Comparison of the size ranges for the landed catches of (a) *Rhynchobatus australiae* using total length (TL cm) and (b) using disk width (DW cm) for *Maculabatis gerrardi, Pateobatis ater, Pateobatis fai, Pateobatis jenkinsii*, and *Pateobatis uarnacoides* caught in the small mesh gillnet (blue), Java Sea trawl fishery (dark green), Arafura Sea trawl fishery (light green), trammel net (purple), hand- and long-line (yellow) and tangle net (red) landed in Muara Angke landing site, Jakarta Indonesia. The size at birth (red dotted line) and size at maturity (black dashed line; M, male; F, female) are displayed from Last et al. (2016).

Similar trends are evident for a number of dasyatid rays, with life stages from neonates to sub-adults w landed in other fisheries during the 2001 – 2005 market surveys (Fig. 6b). Catch of *M. gerrardi* was recorded from small mesh gillnet fishery (neonates – mature adults; ∼20 – 70 cm DW), the Java Sea trawl fishery (neonates – mature adults; ∼20 – 100 cm DW), and the trammel net fishery off southern Java (juveniles – mature adults; ∼30 – 70 cm DW) (Fig. 6b). *Pastinachus ater* was recorded in the hand- and long-line fisheries (∼70 – 110 cm DW), compared to typically larger size range landed in the tangle net fishery (∼80 – 150 cm DW) (Fig. 6b). *Pateobatis fai* was recorded in the Java Sea trawl fishery (juveniles; ∼60 – 70 cm DW), in the Arafura Sea trawl fishery (sub adults; ∼139 – 160 cm DW) and hand- and long-line fisheries (juveniles to mature adults; ∼65 – 160 cm DW) (Fig. 6b). *Pateobatis jenkinsii* was also exposed to fishing throughout all life stages, from small mesh gillnets (neonates – sub adults; ∼25 – 75 cm DW), hand- and long-line fisheries (juveniles – mature adults; ∼40 – 100 cm DW), as well as the tangle net fishery (juveniles – mature adults; ∼60 – 140 cm DW) (Fig. 5b). Catch of *P. uarnacoides* was recorded in the Java Sea trawl fishery (juveniles to mature adults; ∼30 – 60 cm DW), and in the southern Java trammel net fishery (juveniles to mature adults; ∼25 – 55 cm DW) (Fig. 6b).

## 4. Discussion

Wedgefishes and giant guitarfishes face an extremely high risk of extinction (Kyne et al., 2020). While the status of populations and the exact extent of declines for wedgefish and giant guitarfish are uncertain, the current data available suggests that the global population declines are substantial (Kyne et al., 2020, Jabado, 2018, Moore, 2017, Diop and Dossa, 2011). This study advances our understanding of the data poor tangle net fishery in Indonesia, providing details on the species and size composition, selectivity of the fishery for larger size classes, and the consequent declines of shark-like ray populations in Indonesia. In doing so, the study provides important historical context that provides indications of previous, and potentially reference population sizes, the extent of declines, and the shift in target species. Population declines were inferred from the declining catch rates of the target species, *R. australiae*, and the decline in the number of vessels operating in the tangle net fishery. The number of vessels in the fishery peaked at 500 vessels in 1980s and rapidly declining to 100 vessels left fishing in the area in 1996 (Keong, 1996), and further declined to seven vessels in 2017/2018 (see Supplementary Table S5). The significant reduction in the number of vessels operating in the fishery with the marked change in catch composition over the 28 year period, suggests that target species populations have declined and the fishery became economically unviable. Over theses decades, it seems likely that the profits from the fins, salted meat and leather, were insufficient to maintain the tangle net fishery, over the increasing operating costs and decreases in elasmobranch catches (Suzuki, 1997). The tangle net fishery was still operational as of July 2018 (Table S5), and the information collected from this study (i.e. the shift to target stingrays and the values) suggests that the contemporary tangle net fishery is likely reliant on the exploitation of stingrays for its viability and profitability, and opportunistically exploit the high valued wedgefish and guitarfish species when they are landed. However, there is no species-specific catch information for this fishery after 2005 and the landed catch was grouped under single labels such as ‘rays’, ‘sharks’, ‘wedgefish’ and ‘mixed’, and species diversity and catch composition are poorly understood. Given the current state of batoids around the world and the scale of Indonesia’s shark fishery, the contemporary catch of the tangle-net fishery requires further investigation.

Opportunistic exploitation occurs when multiple species can be exploited in the same habitat (Branch et al., 2013). The most desirable and profitable species are targeted and depleted first, before exploitation switches to a less desirable species, leading to overexploitation and harvesting to extinction of both the desired and less desired species (Branch et al., 2013, Burgess et al., 2017). The apparent decline in wedgefish landings and the subsequent increase in stingray landed catches is an indication of the opportunistic exploitation in the fishery. Both groups are common demersal species in the Indo–West Pacific, and occupy similar habitats along inshore continental shelves waters to at least 60 m (White et al., 2006, Compagno and Last, 1999). A decline or change in abundance of wedgefishes in the fishing ground, could result in the stingrays that occupy the same habitat becoming the main target catch, with opportunities to retain the higher valued and critically endangered wedgefish and giant guitarfish species when they are encountered. The time of year may also influence the species catch composition, however more detailed catch information from throughout the year is needed to explore this possibility.

There has been an increasing domestic demand for stingray leather in Indonesia over the past 20 years, with the development of a commercial market for stingray leather on the north coast of Central Java during the period 1980 – 1990 (Ibrahim, 2003). Prior to 1980, stingrays were discard as bycatch in the tangle net fishery (Ibrahim, 2003, Sahubawa et al., 2018), now they are the main component of the landed catch in 2018. The calcified denticles in the stingray skin, such as *P. jenkinsii*, results in a unique and attractive finish to the leather products (e.g. bags, belts, wallets and jewellery), and have a high economic value (Karthikeyan et al., 2009, Sahubawa et al., 2018). Additionally, stingrays remain a major fishery export from Indonesia with stingray meat being consumed throughout the region and conservation concerns growing for species such as *M. gerrardi* and *M. macrura* (Clark-Shen et al., 2021). This targeted fishing pressure on the stingrays may also be resulting in population declines. It is likely that stingrays are experiencing similar trends of declines to shark-like rays, given the increasing pressure on stingrays as wedgefish have become less common. Similar to shark-like rays, there is little fisheries information on tropical stingrays, and their population status is unknown (Smith et al., 2008). The loss of large, benthic elasmobranchs may have significant social, cultural, and economic impacts on the fishers and communities who depend on them (Jaiteh et al., 2017), as well as ecological consequences, altering important ecological processes such as predator-prey interactions and bioturbation from benthic feeding (Kyne and Bennett, 2002, Flowers et al., 2020). The occurrence of large shark-like rays and stingrays and their high economic value has been used as justification for the continuation of the tangle net fishery (Amir, 1988),but the long-term sustainability of the fishery needs to be assessed, especially considering the impacts of other fisheries affecting different size classes.

Length selective fishing mortality often drives a reduction in the catch length composition in heavily fished elasmobranch populations (Walker, 1998, Stevens et al., 2000). At the time of the landing site surveys in 2001 – 2005, a number of the abundant species were landed close to the known maximum sizes. It can thus be inferred that the populations of wedgefishes, giant guitarfishes, and stingrays were experiencing length selective fishing mortality. It is expected that individuals from contemporary populations would be reaching a smaller maximum size and younger maximum age than previous generations due to fishing pressure (Thorson and Simpfendorfer, 2009). Sexual dimorphism is known to occur for numerous elasmobranch species, including *R. australiae, A. ocellatus* and *U. asperrimus* (Last et al., 2016), with females attaining a larger maximum size than males (White and Dharmadi, 2007). Female elasmobranches are more likely to be captured in the large-meshed tangle nets. The majority of *R. australiae, M. gerrardi, and P. uarnacoides* specimens were large females, with 16 individuals of *R. australiae* examined internally being pregnant as reported in White and Dharmadi (2007). The removal of large, breeding individuals from the population causes a reduction in the reproductive potential, resulting in rapid declines in these populations (Prince, 2005). The capture of large breeding females coupled with a sharp decline in catch size suggests that recruitment overfishing may have occurred for several species in the tangle net fishery. Recruitment overfishing occurs when the breeding stock size is reduced to a point where future recruitment declines strongly, and/or the reduction of the number of young entering the population (capture of juveniles), and/or through environmental degradation, which affects the size or suitability of nurseries (Allen et al., 2013, Pauly, 1988, Walters and Martell, 2004). In fisheries where only adults or the juveniles are caught, higher levels of fishing can be sustained (Simpfendorfer, 1999, Prince, 2005).

The impact of other fisheries also likely contributed to these historic declines, and may continue to affect contemporary fisheries. When all age/size classes are fished, it is much more difficult to achieve sustainable outcomes. All life stages of these rays are experiencing fishing mortality from multiple fisheries in Indonesia, as the smaller size classes for many of the species encountered in the tangle net fishery are also caught as bycatch or opportunistic catch in numerous other fisheries operating in Indonesian waters, i.e. trawls, hand- and long-lines, smaller mesh gillnets, and trammel nets. Recruitment overfishing substantially impacts the populations’ productivity and may lead to collapse or prevent the recovery of the population (Allen et al., 2013). Some species of wedgefish and giant guitarfish have a higher than average theoretical population productivity (i.e. recover at a faster rate from population declines) for chondrichthyans (D’Alberto et al., 2019). However, these species still have relatively low reproductive rates and long lifespans compared to teleost fish, and thus can only withstand modest to low levels of fishing mortality (Dulvy et al., 2014b, Cortés, 2000, Camhi et al., 1998, Musick, 1999). Combined with life history information, the magnitude of Indonesia chondrichthyan catches, and the knowledge of the effects of fisheries on large elasmobranch species that mainly takes adults (Prince, 2005, McAuley et al., 2007b, McAuley et al., 2007a), it is likely that these populations of rays are experience unsustainable levels of exploitation and have little potential for recovery without significant reductions in fishing mortality.

The sustainability of fisheries (both harvest and discards) and conservation related research and management initiatives can be compromised by the misidentification of species (Garcia-Vazquez et al., 2012). Incorrect species identification is prevalent throughout multi-species fisheries where visual identification is challenging, and can lead to incidental overfishing, resulting in a higher risk of extinction for the misidentified species in the fishery (Metcalfe et al., 2007). While the latest available trade data from the Food and Agriculture Organization (FAO) does distinguish trade in rays and skates from other elasmobranchs, there is no genera or species specific information (Ramaschiello and Vannuccini, 2015). In South East Asia, *R. australiae* is the most commonly caught wedgefish species (Giles et al., 2016), yet due to similarities in morphology and identification issues, it is commonly confused with other large wedgefish species, in particular with *R. djiddensis, R. laevis* (Giles et al., 2016) and *R. palpebratus* (Compagno and Last, 2008). Historically, all four species of wedgefish have been referred to as ‘white spotted wedgefish’ as the common name and/or *R. djiddensis* as the species name throughout the Indo-Pacific (Last et al., 2016). A recent clarification of species distributions has recognised that *R. djiddensis* is restricted to the Western Indian Ocean (Last et al., 2016). A similar situation has occurred for *G. typus* and *Glaucostegus granulatus*, where there are records of the sharpnose guitarfish *Glaucostegus granulatus* from the tangle net fishery in 1987, where it constituted 4.6% of the total landed catch (Amir, 1988). This is likely to be a misidentification of *G. typus*, as prior to 2016, the range for as *G. granulatus* was poorly described with no records to suggest that this species occurred in Indonesia, and is now known to only occur in the northern Indian Ocean between Myanmar and the Arabian Gulf (Last et al., 2016). Misidentification of elasmobranchs can be further compounded by the ambiguity over the ranges of these species, as some species are rarer in landings and possibly have more of a restricted and/or even fragmented spatial distributions. For example, the broadnose wedgefish *Rhynchobatus springeri* distribution overlaps with *R. australiae* off Java and Sumatra (Giles et al., 2016). The landed catches of small *Rhynchobatus* spp. during the 2001 – 2005 surveys could possibly have included this species, as maximum known size for *R. springeri* is 213 cm TL (Last et al., 2016), compared to 300 cm TL for *R. australiae* (Last et al., 2016, Kyne et al., 2019). The stingrays, *M. gerrardi* and *M. macrura* have overlapping distributions and differ in mostly subtle morphological characteristics (Last et al., 2016), thus without genetic identification, the two cannot be readily differentiated. However, our data could not be retrospectively confirmed as either or both species. As the catch records may comprise both *M. gerrardi* and *M. macrura*, the numbers presented may be an overestimation of *M. gerrardi* catch. This complication in species differentiation is an ongoing issue that complicates efforts to quantify species specific catch and consumption [e.g. Clark-Shen et al. (2021)]. The lack of species specific reporting was thought to masked the known global declines for wedgefish and giant guitarfish throughout their distribution, where from historical and contemporary records, there is evidence of population declines, however the extent of declines are unknown due to the lack of species specific and time-series data (Kyne et al., 2019). Species identification training for fisheries officials and observers, and the use of genetic identification such as in-situ DNA barcoding field kits (Booth et al., 2018), are some methods to address the lack of species specific information and can help with law enforcement in Indonesia and South East Asia (Tillett et al., 2012).

Multi-species fisheries are complex social-ecological systems, and successful management will require significant regulations in governance across local, regional and global scales (Ostrom et al., 1999). As Indonesian government is a signatory party of CITES, it has taken important steps to implement international obligations under CITES and FAO National Plan of Action (NPOA) for the Conservation and Management of Sharks and Rays, with either full species protections (e.g. sawfishes, mobulids) or export controls, and is working to regulate international trade originating from Indonesia. Appendix II listed species are allowed to be landed and traded domestically, and there is no oversight by CITES unless the species is being exported internationally, which requires a positive Non Detriment Finding. These measures would not necessarily affect the harvest and domestic use of these species, and do not cover the other 24 non – CITES elasmobranch species caught in the tangle net fishery. Currently there are no national or regional laws in Indonesia that specifically regulate the take and use of wedgefish, guitarfishes and stingrays (Rusandi et al., 2019). Scientific advice and information (e.g. monitoring of fisheries landings, supply chains) will be required to make management decisions for elasmobranch fisheries in SE Asia (Clark-Shen et al., 2021). Indonesia has complex fisheries supply and trade chains, from fishers to buyers to exporters, which have intricate interactions and different drivers occurring throughout the chain (Rusandi et al., 2019). To ensure the sustainable use of wedgefish, giant guitarfish and stingrays in Indonesia, there will be a need to target the multiple levels of supply chains, from fisheries to exporters, with strategic and evidence based methods (Booth et al., 2020). Improved trade monitoring and traceability, as well as awareness campaigns will be key to reduce fishing mortality, utilisation and trade (Clark-Shen et al., 2021).

Reducing fishing mortality must be directed at managing extraction as well as increasing the compliance and enforcement of trade regulations. For a targeted shark and ray fishery, an example of technical management strategies that could be used to mitigate the risk of overfishing of stingrays, wedgefish and giant guitarfish in the tangle net fishery may involve the use of gear restrictions, and spatial and temporal closures (Harry et al., 2011, Yulianto et al., 2018). Given the size selectivity of the tangle net gear for larger animals, modifying the gear selectivity of the tangle net through the use of smaller mesh gill nets (> 49 cm) will likely result in a case where large and mature individuals for *R. australiae, M. gerrardi*, will rarely be captured and it would likely reduce the fishing pressure on larger size classes of the other rays e.g. *P. ater, P. fai, P. jenkinsii*, and *P. uarnacoides*. Thus adults will be subjected to the ‘gauntlet effect’ and effectively excluded, while juveniles are captured by the fishery. This harvest strategy is considered an effective method to extract long lived species, provided that the fishing mortality on adults (Simpfendorfer, 1999, Prince, 2005) from Indonesian hook and longline fisheries remain low. Research into remediation through post release survivability from different gear types such as trawls and longlines will be beneficial to understand the risks of certain gear types and reducing mortality. Spatial and temporal closures (e.g. closure of nursery grounds) may be used to reduce fishing pressure on these rays, however this will require information on spatial ecology of wedgefish, guitarfish and stingrays (e.g. potential sex/ habitat segregation, locations of nursery areas). However, there is little to no such information presently available for Indonesian waters. It would be difficult to control the level of opportunistic catch and bycatch of rays in all fisheries in Indonesia with limited fisheries management resources and capacity. Local and regional specific management will be required to address the overfishing of stocks, including potential reductions in fishing effort and fishing mortality (Vincent et al., 2014). Any viable management options and regulations will need to ensure that they are leading to noticeable conservation outcomes (Booth et al., 2020), as well as positive social and economic outcomes for fishers (Booth et al., 2019a). Appropriate and economically viable incentives for livelihood alternatives for fishers will be required to address the issues of poverty, food security and lack of alternatives in resources dependent communities (MacKeracher et al., 2020). Failure to do so will result in low compliance and illegal fishing throughout much of their range (Crona et al., 2015, Jaiteh et al., 2017, MacKeracher et al., 2020). Using a precautionary and holistic risk-based approach like a mitigation hierarchy framework as proposed by Booth et al. (2019b) would be highly beneficial in Indonesia as it takes into the account the biological and socio-economic aspect of fisheries. This method has the capability of dealing with data paucity, and could be used for the tangle net fishery. Regional management across multiple fisheries and life stages will be required to ensure the sustainability and conservation of wedgefish, giant guitarfish and stingrays that are being caught across a wide range of fisheries at all life stages.

## 5. Conclusion

Evidence of substantial and rapid declines in landings of the target species raises concerns about the status of shark-like rays in Indonesia. The tangle net fishery was established to target wedgefish in the 1980s due to the high value of the fins, but evidence suggests populations of wedgefish have declined dramatically in many areas. The tangle net fishery remained active in 2018, however the fishery now appears to be mainly catching large stingrays, which are used for leather and meat, with wedgefishes only occasionally caught. The continuing pressure on stingrays may also be resulting in population declines, and the extent of such declines is unknown. Some ray species landed in the tangle net fishery were also caught in other fisheries at different life stages, requiring management across multiple fisheries and life stages, instead of single fishery management, to ensure the sustainability and conservation of rays. There are concerns about the sustainability of the fishery and the population status of many of ray species targeted. The results of this study emphasise the urgency for effective management for the conservation of these rays.

## Acknowledgements

The authors would like to thank the staff of the fish auction office for the Indonesian Centre for Fisheries Research for the collection of the landed catch vessels at the Muara Angke landing port during July 2016 – 2018. This work was funded by the Shark Conservation Fund, a philanthropic collaborative pooling expertise and resources to meet the threats facing the world’s sharks and rays (B.M.D.). The Shark Conservation Fund is a project of Rockefeller Philanthropy Advisors. The landing port surveys between 2001 and 2005 were funded by the Australian Centre for International Agricultural Research (ACIAR; grants FIS/2000/062 and FIS/2003/037) and CSIRO Oceans & Atmosphere (W.T.W.). B.M.D. was supported through an Australian Government Research Training Program Scholarship (RTPS). The funders had no role in study design, data collection and analysis, decision to publish, or preparation of the manuscript. The authors would like to thank Dr Stephanie Hernandez for the design of the map. Lastly, we dedicate this paper to the memory of ‘Pak’ Dharmadi, a leader and pioneer in shark and rays research and management in Indonesia.

## Contributions

The manuscript was conceptualised by W.T.W., B.M.D., and C.A.S. The data was collected by W.T.W. and D.D., and analysed by B.M.D., and W.T.W. The manuscript was written by B.M. D., which was reviewed and edited by W.T.W., A.C., D.D., and C.A.S.

## Competing interests

The authors declare no competing interests.

## Data availability

All data generated or analysed during this study are included in this published article (and its Supplementary Information files).

## Notes

### Competing Interest Statement

The authors have declared no competing interest.

