## Supplemental Table for "Untangling the Indonesian tangle net fishery: describing a data-poor fishery targeting large threatened rays (Order Batoidea)"

**Table S1.** Dates of the landing port surveys conducted in the Muara Angke landing port, and the associated village processing area in Jakarta April 2001 - December 2005. Presence or absence of tangle net vessels at the landing port as recorded (Yes or No). The total number of species and specimens for rays and sharks per survey day are reported.

| **Date** | **Muara Angke** | **Village processing area** | **No. of rays species** | **No. of ray specimens** | **No. of shark species** | | **No. of shark specimens** |
| --- | --- | --- | --- | --- | --- | --- | --- |
| 4/04/2001 | **Yes** | No | 14 | Unknown | 3 | | Unknown |
| 5/04/2001 | **Yes** | No | 15 | 11* | 2 | | Unknown |
| 6/04/2001 | No | No | ­­ ­­ | ­­ ­­ | ­­ ­­ | | ­­ ­­ |
| 27/06/2001 | No | No | ­­ ­­ | ­­ ­­ | ­­ ­­ | | ­­ ­­ |
| 28/06/2001 | No | No | ­­ ­­ | ­­ ­­ | ­­ ­­ | | ­­ ­­ |
| 11/07/2001 | No | No | ­­ ­­ | ­­ ­­ | ­­ ­­ | | ­­ ­­ |
| 13/03/2002 | No | No | ­­ ­­ | ­­ ­­ | ­­ ­­ | | ­­ ­­ |
| 14/03/2002 | No | No | ­­ ­­ | ­­ ­­ | ­­ ­­ | | ­­ ­­ |
| 17/03/2002 | No | No | ­­ ­­ | ­­ ­­ | ­­ ­­ | | ­­ ­­ |
| 20/03/2002 | No | No | ­­ ­­ | ­­ ­­ | ­­ ­­ | | ­­ ­­ |
| 15/05/2002 | **Yes** | No | 10 | 159 | 0 | | 0 |
| 16/05/2002 | No | No | ­­ ­­ | ­­ ­­ | ­­ ­­ | | ­­ ­­ |
| 19/05/2002 | No | No | ­­ ­­ | ­­ ­­ | ­­ ­­ | | ­­ ­­ |
| 20/05/2002 | No | No | ­­ ­­ | ­­ ­­ | ­­ ­­ | | ­­ ­­ |
| 14/08/2002 | No | **Yes** | 15 | 197 | 2 | | 3 |
| 15/08/2002 | No | No | ­­ ­­ | ­­ ­­ | ­­ ­­ | | ­­ ­­ |
| 18/08/2002 | No | No | ­­ ­­ | ­­ ­­ | ­­ ­­ | | ­­ ­­ |
| 18/10/2002 | **Yes** | No | 9 | 96 | 0 | | 0 |
| 19/10/2002 | No | No | ­­ ­­ | ­­ ­­ | ­­ ­­ | | ­­ ­­ |
| 31/01/2003 | No | No | ­­ ­­ | ­­ ­­ | ­­ ­­ | | ­­ ­­ |
| 1/02/2003 | No | No | ­­ ­­ | ­­ ­­ | ­­ ­­ | | ­­ ­­ |
| 9/02/2003 | No | **Yes** | 10 | 361 | 0 | | 0 |
| 19/04/2004 | No | No | ­­ ­­ | ­­ ­­ | ­­ ­­ | | ­­ ­­ |
| 21/04/2004 | No | **Yes˟** | ­­ ­­ | ­­ ­­ | ­­ ­­ | | ­­ ­­ |
| 22/04/2004 | No | No | ­­ ­­ | ­­ ­­ | ­­ ­­ | | ­­ ­­ |
| 17/07/2004 | **Yes** | No | 13 | 147 | 2 | | 4 |
| 18/07/2004 | No | No | ­­ ­­ | ­­ ­­ | ­­ ­­ | | ­­ ­­ |
| 20/07/2004 | No | **Yes** | 1 | 49 | 0 | | 0 |
| 21/07/2004 | **Yes** | No | 13 | 131 | 2 | | 3 |
| 30/08/2004 | No | No | ­­ ­­ | ­­ ­­ | ­­ ­­ | | ­­ ­­ |
| 31/08/2004 | No | No | ­­ ­­ | ­­ ­­ | ­­ ­­ | | ­­ ­­ |
| 1/09/2004 | No | No | ­­ ­­ | ­­ ­­ | ­­ ­­ | | ­­ ­­ |
| 4/10/2004 | No | No | ­­ ­­ | ­­ ­­ | ­­ ­­ | | ­­ ­­ |
| 5/10/2004 | No | No | ­­ ­­ | ­­ ­­ | ­­ ­­ | | ­­ ­­ |
| 6/10/2004 | No | **Yes** | 12 | 106 | 4 | | 5 |
| 6/12/2004 | No | No | ­­ ­­ | ­­ ­­ | ­­ ­­ | | ­­ ­­ |
| 7/12/2004 | No | No | ­­ ­­ | ­­ ­­ | ­­ ­­ | | ­­ ­­ |
| 8/12/2004 | No | No | ­­ ­­ | ­­ ­­ | ­­ ­­ | | ­­ ­­ |
| 23/01/2005 | No | No | ­­ ­­ | ­­ ­­ | ­­ ­­ | | ­­ ­­ |
| 24/01/2005 | No | No | ­­ ­­ | ­­ ­­ | ­­ ­­ | | ­­ ­­ |
| 13/03/2005 | No | No | ­­ ­­ | ­­ ­­ | ­­ ­­ | | ­­ ­­ |
| 14/03/2005 | No | No | ­­ ­­ | ­­ ­­ | ­­ ­­ | | ­­ ­­ |
| 15/03/2005 | No | No | ­­ ­­ | ­­ ­­ | ­­ ­­ | | ­­ ­­ |
| 31/05/2005 | No | No | ­­ ­­ | ­­ ­­ | ­­ ­­ | | ­­ ­­ |
| 1/06/2005 | No | No | ­­ ­­ | ­­ ­­ | ­­ ­­ | | ­­ ­­ |
| 2/06/2005 | No | No | ­­ ­­ | ­­ ­­ | ­­ ­­ | | ­­ ­­ |
| 15/07/2005 | No | No | ­­ ­­ | ­­ ­­ | ­­ ­­ | | ­­ ­­ |
| 16/07/2005 | **Yes** | No | 8 | 51 | 0 | | 0 |
| 4/10/2005 | No | No | ­­ ­­ | ­­ ­­ | ­­ ­­ | | ­­ ­­ |
| 5/10/2005 | **Yes** | **Yes** | 14 | 239 | 5 | | 8 |
| 7/10/2005 | No | No | ­­ ­­ | ­­ ­­ | ­­ ­­ | | ­­ ­­ |
| 7/12/2005 | No | No | ­­ ­­ | ­­ ­­ | ­­ ­­ | | ­­ ­­ |
| 8/12/2005 | No | **Yes** | 1 | 4 | 1 | | 2 |
| **Total** | 8 | 7 | ­­ | 1534 | ­­ | | 25 |
| * Only 11 specimens were recorded from 5 species of rays, no information on the remaining 6 species of rays and sharks were documented | | | | | | | |
| **˟** No specimens were recorded on this day, informal interviews were conducted with the fishers on the price of fins, meat and skins | | | | | |  | |

**Table S2.** Length-weight relationship for the elasmobranch species caught in the Indonesian tangle-net fishery and landed in Muara Angke, Jakarta April 2001–December 2005.

| **Family** | **Species** | ***a*** | ***b*** | ***R*^2^** | **Source** |
| --- | --- | --- | --- | --- | --- |
| Pristidae | *Pristis pristis* | 0.003 | 2.9985 | 0.949 | Salini et al. (2007) |
| Glaucostegidae | *Glaucostegus thouini* | 0.0046 | 2.9184 | 0.9797 | White (2018) |
|  | *Glaucostegus typus* | 0.0046 | 2.9184 | 0.9797 | White (2018) |
| Rhinidae | *Rhina ancylostoma* | 0.008 | 3.012 | 0.9988 | from length and weights in Gordon (1992); Rajapackiam et al. (2007); Wallace (1967) |
|  | *Rhynchobatus australiae* | 0.0045 | 2.9959 | 0.987 | For *R. palpebratus*  White et al. (2019) |
|  | *Rhynchobatus palpebratus* | 0.0045 | 2.9959 | 0.987 | White et al. (2019) |
| Dasyatidae | *Bathytoshia lata* | -- | -- | -- | Weight from Struhsaker (1969) |
|  | *Himantura leoparda* | 0.0728 | 2.7578 | 0.9737 | For *H. australis*  White et al. (2019) |
|  | *Himantura uarnak* | 0.0728 | 2.7578 | 0.9737 | For *H. australis*  White et al. (2019) |
|  | *Himantura undulata* | 0.0728 | 2.7578 | 0.9737 | For *H. australis*  White et al. (2019) |
|  | *Maculabatis astra* | 0.0219 | 3.0471 | 0.9844 | White et al. (2019) |
|  | *Maculabatis gerrardi* | 0.0219 | 3.0471 | 0.9844 | White et al. (2019) |
|  | *Megatrygon microps* | 0.0728 | 2.7578 | 0.9737 | For *H. australis*  White et al. (2019) |
|  | *Pastinachus ater* | 0.0728 | 2.7578 | 0.9737 | For *H. australis*  White et al. (2019) |
|  | *Pateobatis fai* | 0.0728 | 2.7578 | 0.9737 | For *H. australis*  White et al. (2019) |
|  | *Pateobatis jenkinsii* | 0.0728 | 2.7578 | 0.9737 | For *H. australis*  White et al. (2019) |
|  | *Pateobatis uarnacoides* | 0.0728 | 2.7578 | 0.9737 | For *H. australis*  White et al. (2019) |
|  | *Taeniurops meyeni* | 0.0728 | 2.7578 | 0.9737 | For *H. australis*  White et al. (2019) |
|  | *Urogymnus asperrimus* | 0.0728 | 2.7578 | 0.9737 | For *H. australis*  White et al. (2019) |
|  | *Urogymnus granulatus* | 0.0728 | 2.7578 | 0.9737 | For *H. australis*  White et al. (2019) |
| Gymnuridae | *Gymnura zonura* | 0.0050 | 3.078 | 0.977 | White and Dharmadi (2007) |
| Aetobatidae | *Aetobatus ocellatus* | 0.0276 | 2.87 | 0.98 | for *A. ocellatus/narinari*  Bassos-Hull et al. (2014) |
| Myliobatidae | *Aetomylaeus vespertilio* | 0.0276 | 2.87 | 0.98 | for *A. ocellatus/narinari*  Bassos-Hull et al. (2014) |
| Carcharhinidae | *Carcharhinus amboinensis* | 0.00194 | 3.27 | 0.986 | Stevens and McLoughlin (1991) |
|  | *Carcharhinus obscurus* | 0.000032 | 2.7862 | 0.9649 | Kohler et al. (1996) |
|  | *Carcharhinus leucas* | 0.0111 | 2.923 | 0.908 | Froese et al. (2014); Compagno (1984) |
|  | *Carcharhinus limbatus* | 0.00251 | 3.125 | 0.989 | Castro (1996) |
| Galeocerdidae | *Galeocerdo cuvier* | 0.00141 | 3.24 | -- | Randall (1992) |
| Ginglymostomatidae | *Nebrius ferrugineus* | 0.009006 | 2.911 | 0.988 | Castro (2000) |
| Hemigaleidae | *Hemipristis elongatus* | 0.00162 | 3.21 | -- | Stevens and McLoughlin (1991) |
| Sphyrnidae | *Sphyrna lewini* | 0.00399 | 3.03 | 0.985 | Stevens and Lyle (1989) |
|  | *Sphyrna mokarran* | 0.00123 | 3.24 | 0.991 | Stevens and Lyle (1989) |

**Table S3.** Temporal occurrence of elasmobranchs landed from tangle net vessels in Muara Angke landing site, Jakarta in April 2001–December 2005, over a total of 53 survey days, grouped by months that were surveyed.

|  |  | **2001** | | | **2002** | | | | **2003** | | **2004** | | | | | | | **2005** | | | | | | |
| --- | --- | --- | --- | --- | --- | --- | --- | --- | --- | --- | --- | --- | --- | --- | --- | --- | --- | --- | --- | --- | --- | --- | --- | --- |
| **Family** | **Species** | Apr | Aug | Jul | Mar | May | Aug | Oct | Jan | Feb | Apr | Jul | Aug | Sep | Oct | Dec | Jan | | Mar | May | Jun | Jul | Oct | Dec |
| Pristidae | *Pristis pristis* | -- | -- | -- | -- | -- | Y | -- | -- | -- | -- | -- | -- | -- | -- | -- | -- | | -- | -- | -- | -- | -- | -- |
| Glaucostegidae | *Glaucostegus thouini* | Y | -- | -- | Y | -- | -- | -- | -- | -- | -- | -- | -- | -- | -- | -- | -- | | -- | -- | -- | -- | -- | -- |
|  | *Glaucostegus typus* | Y | -- | -- | -- | -- | -- | -- | -- | -- | -- | Y | -- | -- | -- | -- | -- | | -- | -- | -- | Y | -- | -- |
| Rhinidae | *Rhina ancylostoma* | Y | -- | -- | -- | -- | -- | Y | -- | -- | -- | Y | -- | -- | Y | -- | -- | | -- | -- | -- | Y | Y | Y |
|  | *Rhynchobatus australiae* | Y | -- | -- | Y | -- | Y | Y | -- | -- | -- | Y | -- | -- | Y | -- | -- | | -- | -- | -- | Y | Y | -- |
|  | *Rhynchobatus palpebratus* | -- | -- | -- | -- | -- | Y | -- | -- | -- | -- | -- | -- | -- | -- | -- | -- | | -- | -- | -- | -- | -- | -- |
| Dasyatidae | *Bathytoshia lata* | -- | -- | -- | -- | -- | Y | -- | -- | -- | -- | -- | -- | -- | -- | -- | -- | | -- | -- | -- | -- | -- | -- |
|  | *Himantura leoparda* | Y | -- | -- | -- | -- | Y | -- | -- | Y | -- | Y | -- | -- | Y | -- | -- | | -- | -- | -- | Y | Y | -- |
|  | *Himantura uarnak* | Y | -- | -- | Y | -- | Y | -- | -- | Y | -- | Y | -- | -- | Y | -- | -- | | -- | -- | -- | -- | Y | -- |
|  | *Himantura undulata* | Y | -- | -- | -- | -- | -- | -- | -- | -- | -- | -- | -- | -- | -- | -- | -- | | -- | -- | -- | -- | Y | -- |
|  | *Maculabatis astra* | -- | -- | -- | -- | -- | Y | -- | -- | -- | -- | -- | -- | -- | -- | -- | -- | | -- | -- | -- | -- | -- | -- |
|  | *Maculabatis gerrardi* | Y | -- | -- | Y | -- | -- | Y | -- | Y | -- | Y | -- | -- | Y | -- | -- | | -- | -- | -- | Y | Y | -- |
|  | *Megatrygon microps* | -- | -- | -- | -- | -- | -- | -- | -- | -- | -- | Y | -- | -- | -- | -- | -- | | -- | -- | -- | -- | -- | -- |
|  | *Pastinachus ater* | Y | -- | -- | Y | -- | Y | Y | -- | -- | -- | Y | -- | -- | Y | -- | -- | | -- | -- | -- | Y | Y | -- |
|  | *Pateobatis fai* | Y | -- | -- | Y | -- | Y | Y | -- | Y | -- | Y | -- | -- | Y | -- | -- | | -- | -- | -- | -- | Y | -- |
|  | *Pateobatis jenkinsii* | Y | -- | -- | Y | -- | Y | Y | -- | -- | -- | Y | -- | -- | Y | -- | -- | | -- | -- | -- | Y | Y | -- |
|  | *Pateobatis uarnacoides* | Y | -- | -- | -- | -- | Y | Y | -- | Y | -- | Y | -- | -- | Y | -- | -- | | -- | -- | -- | -- | Y | -- |
|  | *Taeniurops meyeni* | Y | -- | -- | Y | -- | Y | -- | -- | Y | -- | Y | -- | -- | Y | -- | -- | | -- | -- | -- | -- | Y | -- |
|  | *Urogymnus asperrimus* | Y | -- | -- | -- | -- | Y | -- | -- | -- | -- | -- | -- | -- | -- | -- | -- | | -- | -- | -- | -- | -- | -- |
|  | *Urogymnus granulatus* | Y | -- | -- | Y | -- | Y | -- | -- | Y | -- | Y | -- | -- | Y | -- | -- | | -- | -- | -- | -- | Y | -- |
| Gymnuridae | *Gymnura zonura* | Y | -- | -- | -- | -- | -- | Y | -- | Y | -- | Y | -- | -- | -- | -- | -- | | -- | -- | -- | -- | Y | -- |
| Aetobatidae | *Aetobatus ocellatus* | Y | -- | -- | Y | -- | Y | Y | -- | -- | -- | Y | -- | -- | Y | -- | -- | | -- | -- | -- | Y | Y | -- |
| Myliobatidae | *Aetomylaeus vespertilio* | Y | -- | -- | -- | -- | -- | -- | -- | Y | -- | Y | -- | -- | Y | -- | -- | | -- | -- | -- | -- | -- | -- |
| Carcharhinidae | *Carcharhinus amboinensis* | Y | -- | -- | -- | -- | -- | -- | -- | -- | -- | -- | -- | -- | -- | -- | -- | | -- | -- | -- | -- | -- | -- |
|  | *Carcharhinus obscurus* | -- | -- | -- | -- | -- | Y | -- | -- | -- | -- | -- | -- | -- | -- | -- | -- | | -- | -- | -- | -- | Y | -- |
|  | *Carcharhinus leucas* | -- | -- | -- | -- | -- | -- | -- | -- | -- | -- | Y | -- | -- | -- | -- | -- | | -- | -- | -- | -- | Y | -- |
|  | *Carcharhinus limbatus* | Y | -- | -- | -- | -- | Y | -- | -- | -- | -- | Y | -- | -- | Y | -- | -- | | -- | -- | -- | -- | Y | -- |
|  | *Galeocerdo cuvier* | Y | -- | -- | -- | -- | Y | -- | -- | -- | -- | -- | -- | -- | -- | -- | -- | | -- | -- | -- | -- | Y | -- |
| Ginglymostomatidae | *Nebrius ferrugineus* | -- | -- | -- | -- | -- | -- | -- | -- | -- | -- | -- | -- | -- | -- | -- | -- | | -- | -- | -- | -- | Y | -- |
| Hemigaleidae | *Hemipristis elongatus* | -- | -- | -- | -- | -- | -- | -- | -- | -- | -- | -- | -- | -- | Y | -- | -- | | -- | -- | -- | -- | -- | Y |
| Sphyrnidae | *Sphyrna lewini* | Y | -- | -- | -- | -- | -- | -- | -- | -- | -- | Y | -- | -- | Y | -- | -- | | -- | -- | -- | -- | -- | -- |
|  | *Sphyrna mokarran* | Y | -- | -- | -- | -- | -- | -- | -- | -- | -- | Y | -- | -- | Y | -- | -- | | -- | -- | -- | -- | -- | -- |
|  | Number of ray species | 18 | 0 | 0 | 10 | 0 | 14 | 9 | 0 | 9 | 0 | 16 | 0 | 0 | 13 | 0 | 0 | | 0 | 0 | 0 | 8 | 14 | 1 |
|  | Number of shark species | 5 | 0 | 0 | 0 | 0 | 3 | 0 | 0 | 0 | 0 | 4 | 0 | 0 | 4 | 0 | 0 | | 0 | 0 | 0 | 0 | 5 | 1 |
| **Total number of species observed** | | 23 | 0 | 0 | 10 | 0 | 17 | 9 | 0 | 9 | 0 | 20 | 0 | 0 | 17 | 0 | 0 | | 0 | 0 | 0 | 8 | 19 | 2 |

**Table S4.** Size at maturity (DW/TL cm) for species with length frequency data in the Indonesian tangle-net fishery and landed in Muara Angke, Jakarta April 2001–December 2005.

| **Species** | **Females** | **Males** | **Source** |
| --- | --- | --- | --- |
| *Rhynchobatus australiae* | – | 131 | Compagno and Last (1999); White (2018) |
| *Himantura leoparda* | – | 70 – 94 | White and Dharmadi (2007); Last and Stevens (2009) |
| *Himantura uarnak* | – | 82-84 | White and Dharmadi (2007); White et al. (2006); Manjaji (2004) |
| *Maculabatis gerrardi* | 54 | 48 | White and Dharmadi (2007); White et al. (2006); Manjaji (2004) |
| *Pastinachus ater* | – | 103 | White (2018) |
| *Pateobatis fai* | – | 108 – 122 | White and Dharmadi (2007); Last and Stevens (2009) |
| *Pateobatis jenkinsii* | – | 75 – 85 | White and Dharmadi (2007); Last and Stevens (2009) |
| *Pateobatis uarnacoides* | – | 76 | White et al. (2006); Last and Compagno (1999) |
| *Taeniurops meyeni* | – | 100 – 110 | White (2018) |
| *Aetobatus ocellatus* | 100-110 | 130 | Schluessel et al. (2010); Last et al. (2016) |

**Contemporary Muara Angke tangle net fishery: 2017 – 2018**

**Methods**

Contemporary information on the tangle net fishery from the Muara Angke landing survey data were obtained from the Indonesian Centre for Fisheries Research, from 2^nd^ January 2017 – 16^th^ July 2018. The landed catch from every vessel, including vessels with no catch that landed at Muara Angke site was recorded in Bahasa Indonesia by the staff of the fish auction office for the Indonesian Centre for Fisheries Research. The date of arrival into the landing port, vessel, owner, fishing gear, total number of animals caught, and the main three species and landed catch caught per species in kilograms (kg) was recorded for each vessel. The data was translated to English by Dharmadi in 2018.

**Results**

A total of 198 vessel landings were recorded in the Muara Angke landing site during 2^nd^ January 2017 to 16^th^ July 2018. Of these landings, 14 were tangle net vessel landings from seven individual tangle net vessels (Table 4). The species are only recorded under single labels in Bahasa Indonesia of “yong bung/cucut liong bung” [=wedgefish/shark ray], “pari” [=rays], “cucut” [=sharks], “manyung” [=local catfish species, *Netuma* spp.] and ‘mix/mixed species’. Unidentified wedgefish species comprised a small component of the total landings for these tangle net vessels, with an estimated landed catch of 6 tonnes (Table 4). The majority of catch was recorded as rays with estimated landed catch of 43.9 tonnes, and unknown shark species were recorded once with 200 kg (Table 4). The local catfish

species (*Netuma spp*.) was recorded from the tangle net landings, with an estimated catch of 5 tonnes, while unknown ‘mixed species’ accounted for 8 tonnes (Table 4). No other data on species composition was recorded.

**Table S5.** Indonesian Capture Fisheries data on seven vessel landings from the tangle net fishery recorded at Muara Angke port in June 2017 – July 2018. The date of landing, vessel identifier (vessel ID) fishing gear, with main catch species, and landed catch weight (kilograms, kg) are reported. Dashed lines indicated no data was recorded. The species were recorded in Bahasa Indonesia, here they are reported in English with translation from Dharmadi. Source: Centre For Fisheries Research (2018)

| **Date of landing** | **Vessel ID** | **Species 1** | **Catch (kg)** | **Species 2** | **Catch (kg)** | **Species 3** | **Catch (kg)** | **Total** |
| --- | --- | --- | --- | --- | --- | --- | --- | --- |
| 04/06/2017 | SC-DA-040617 | Rays | 12,000 | -- | -- | -- | -- | 12,000 |
| 28/07/2017 | HJ-OG-280717 | -- | -- | -- | -- | -- | -- | -- |
| 13/09/2017 | HJ-OG-130917 | Rays | 6,000 | Mixed species | 2,500 | -- | -- | 8,500 |
| 03/11/2017 | BH-DA-031117 | Rays | 6,500 | Mixed species | 3,000 | -- | -- | 9,500 |
| 24/11/2017 | KE-OG-241117 | Rays | 3,500 | *Netuma spp.* | 1,000 | Mixed species | 2,000 | 6,500 |
| 07/12/2017 | KE-OG-071217 | -- | -- | -- | -- | -- | -- | -- |
| 11/01/2018 | KE-OG-110118 | -- | -- | -- | -- | -- | -- | -- |
| 29/01/2018 | HJ-OG-290118 | Rays | 4,500 | Wedgefish spp 1 | 1,000 | *Netuma spp.* | 1,000 | 6,500 |
| 13/03/2018 | BH-DA-130318 | Rays | 20,000 | -- | -- |  |  | 20,000 |
| 20/03/2018 | KE-OG-200318 | Sharks | 200 | *Netuma spp.* | 1,000 | Rays | 3,000 | 4,200 |
| 29/03/2018 | TS-HS-290318 | -- | -- | -- | -- | -- | -- | -- |
| 06/06/2018 | SC-DA-060618 | -- | -- | -- | -- | -- | -- | -- |
| 26/06/2018 | KE-OG-260618 | Rays | 5,900 | *Netuma spp.* | 2,000 | -- | -- | 7,900 |
| 07/07/2018 | TS-HS-070718 | Rays | 2,503 | Wedgefish spp 2 | 5,000 | -- | -- | 7,503 |
